# Design of synthetic human gut microbiome assembly and function

**DOI:** 10.1101/2020.08.19.241315

**Authors:** Ryan L. Clark, Bryce M. Connors, David M. Stevenson, Susan E. Hromada, Joshua J. Hamilton, Daniel Amador-Noguez, Ophelia S. Venturelli

**Affiliations:** Department of Biochemistry, University of Wisconsin-Madison, Madison, WI 53706; Department of Bacteriology, University of Wisconsin-Madison, Madison, WI 53706; Department of Chemical & Biological Engineering, University of Wisconsin-Madison, Madison, WI 53706

## Abstract

The assembly of microbial communities and functions emerge from a complex and dynamic web of interactions. A major challenge in microbiome engineering is identifying organism configurations with community-level behaviors that achieve a desired function. The number of possible subcommunities scales exponentially with the number of species in a system, creating a vast experimental design space that is challenging to even sparsely traverse. We develop a model-guided experimental design framework for microbial communities and apply this method to explore the functional landscape of the health-relevant metabolite butyrate using a 25-member synthetic human gut microbiome community. Based on limited experimental measurements, our model accurately forecasts community assembly and butyrate production at every possible level of complexity. Our results elucidate key ecological and molecular mechanisms driving butyrate production including inter-species interactions, pH and hydrogen sulfide. Our model-guided iterative approach provides a flexible framework for understanding and predicting community functions for a broad range of applications.

## INTRODUCTION

Microbial communities carry out pivotal chemical transformations in nearly every environment on Earth^1^. Many of these processes critically impact human health and environmental sustainability, including oceanic CO_2_-fixation^2^, production of growth-promoting molecules in the plant rhizosphere^3^, and degradation of indigestible dietary substrates^4^. Microbial community dynamics and functions are determined by complex and dynamic interactions between constituent community members and their environment. Developing the capabilities to engineer microbiome properties holds promise to address grand challenges facing human society^5^ and methods to predict microbiome functions are needed to enable microbiome engineering efforts.

A bottom-up approach to build and characterize synthetic microcosms has key advantages including reduced complexity compared to natural systems, ability to manipulate environmental parameters and community membership and achieve a high temporal resolution. Previous studies have leveraged synthetic microcosms of bacteria isolated from the human gut^6^ or soil^7^ to demonstrate that dynamic models based on pairwise interactions are predictive of multi-species community assembly. In addition, modeling community assembly using pairwise interactions has provided a deeper understanding of the effects of environmental factors including pH^8^, dilution^9^, nutrient availability^10^, toxins^11^, and temperature^12^ on microbial community behaviors.

A key challenge for predicting microbiome properties is mapping community composition to community-level metabolic functions. Genome-scale metabolic models have been used to predict collective metabolic outputs of microbial communities, an approach which is limited by the quality of functional gene annotations and stringent assumptions^13^. Bottom-up assembly of microbial consortia coupled to mathematical modeling has been used to interrogate how the production or consumption of molecules changes in a community context relative to individual species^11,14,15^. However, computational frameworks to predict both community dynamics and functional outputs for high-dimensional communities that mirror the complexity of natural microbiomes are needed to harness the potential of microbiome engineering for diverse applications.

A detailed and quantitative understanding of microbial interaction networks would enable the design of microbial consortia with robust target functions from the bottom up. While model-guided design^16^ has been used to identify gut microbial communities that elicit a target immune response in mouse models, the complexity of the host system and the low-throughput of mouse studies limits the observability of system parameters and a comprehensive understanding of ecological factors shaping microbiome behaviors. Here, we use a data-driven approach to build a model of butyrate production by complex *in vitro* communities of human-associated intestinal isolates. Butyrate production is a major function of the gut microbiome associated with protection from a wide range of human diseases, including arthritis^17^, diet-induced obesity^18–20^, colitis^21,22^, opportunistic pathogen infection^23^, diabetes^24^, and colorectal cancer^25^. Our approach leverages data-driven models to quantify interactions impacting the growth dynamics of functional organisms and interpretable statistical models to quantify interactions impacting metabolic activities (functional yield of butyrate per unit biomass). By modeling these two interaction types separately, we demonstrate that in some contexts, accurate prediction of functional organism abundance can predict function, while in other community contexts containing metabolically flexible ecological driver species, interactions modifying metabolic modes must be captured to predict function. We use these models to design communities of up to 25 species with a broad range of butyrate production capabilities and analyze our model as well as the metabolic profiles and environmental modification of designed consortia to provide key insights into metabolic interactions impacting butyrate production.

## RESULTS

Identifying highly functional microbial communities from the bottom-up is a major challenge because the number of sub-communities exponentially increases with the system dimension^26^. To explore community design space, we develop a modeling framework to guide iterative design of experiments (**Figure 1a,b**). Ecosystem functions can be modulated by selection effects, defined as changes in function correlated with changes in the abundance of functional species, or complementarity effects, defined as changes in the functional yield per unit biomass for each functional species^27–29^. We implement a dual module modeling framework to determine the contributions of microbe-microbe interactions to each of these effect types. A community dynamic model, referred to as the generalized Lotka-Volterra model (gLV), predicts community assembly and the function model predicts a functional activity from community composition (**Figure 1a**). The gLV model is an ordinary differential equation model that captures the temporal change in species abundances due to monospecies growth parameters and inter-species interactions and has been used to predict and analyze multi-species community assembly based on measurements of lower-order communities^6^. Our function model consists of a regression model with interaction terms mapping species abundance at a specific time point to the concentration of an output metabolite. The inter-species interaction terms in the gLV model represent selection effects (i.e. how one species impacts the growth of another) and the interaction terms in the regression model represent complementarity effects (i.e. deviations from constant yield of a function per unit biomass^15^) (**Figure 1a,c**). For the gLV model, we use Bayesian parameter inference techniques to determine the uncertainty in our parameters based on biological and technical variability in the experimental data^30^. The composite gLV and statistical models predict the probability distribution of the functional activity given an initial condition of species abundances (**Figure 1a, Methods**).

**Figure 1.**
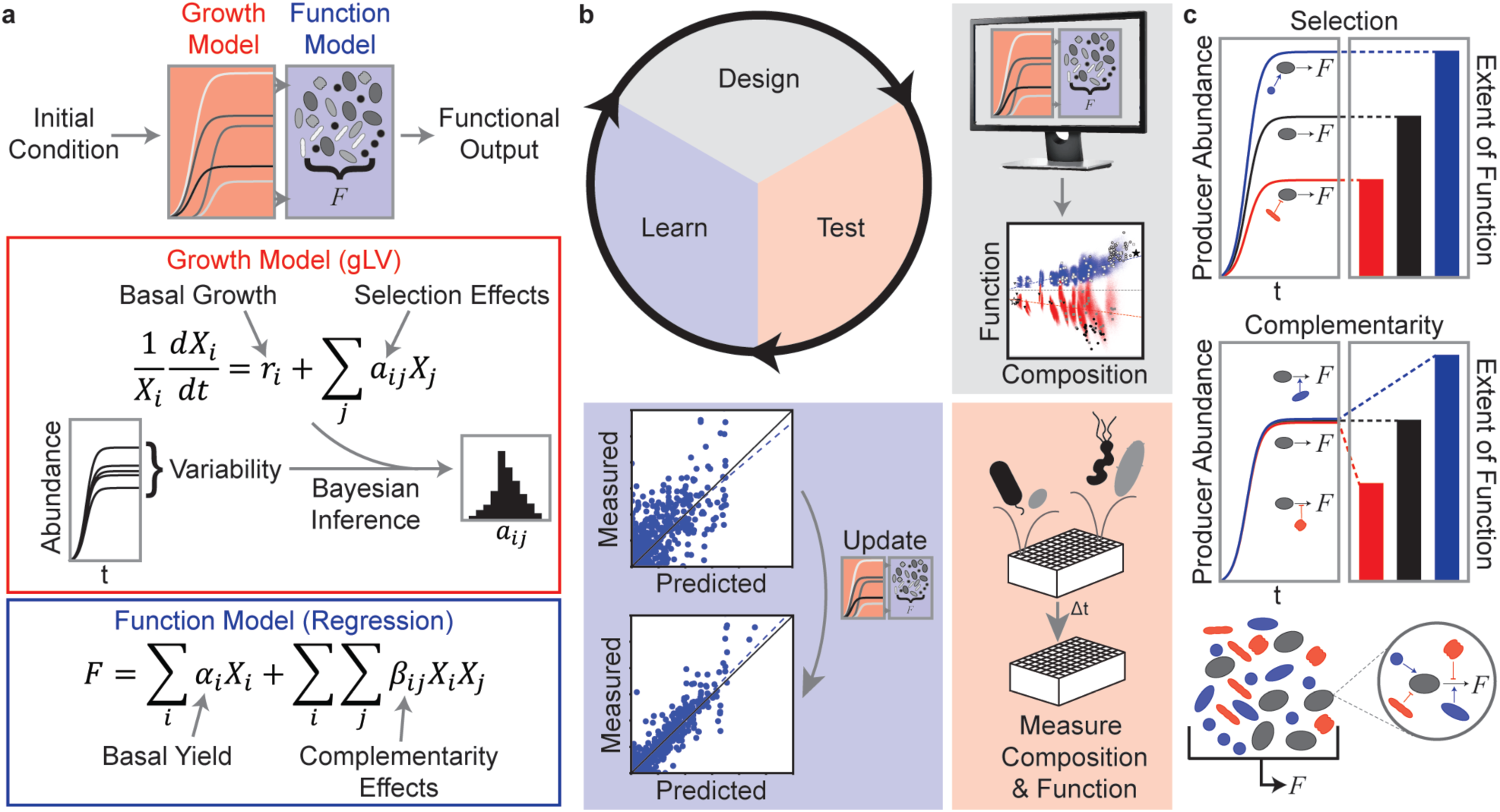
Iterative modeling framework to predict microbial community assembly and function. **(a)** Two-stage modeling framework for predicting community assembly and function. The generalized Lotka-Volterra model (gLV) represents community dynamics. The inter-species interaction terms represent the selection effects described in **c**. A Bayesian Inference approach was used to determine parameter uncertainties due to biological and technical variability. A linear regression model with interactions represents the complementarity effects as described in **c**. Combining these two models enables prediction of a probability distribution of the functional activity from initial species concentrations. **(b)** Model-guided iterative experimental approach for developing a model to predict community assembly and butyrate production. First, we use our model to explore the design space of possible experiments (i.e. different initial conditions of species presence/absence) and design communities that span the range of expected functional outputs. Next, we use high-throughput experimental methods to measure species abundance and functional outputs. Finally, we evaluate the model’s capability to accurately predict the experimental data and train the model on new data for the next iteration. **(c)** Inter-species interactions that impact the functional output of an organism can be driven by selection (top) or complementarity (bottom) effects. In this model, the total functional output of the communities is determined by a combination of these effects.

Due to significant interest in development of defined bacterial therapeutics for human health applications^31^ and the beneficial role of butyrate produced by gut microbiota on a myriad of health outcomes^17–25,32^, we sought to apply our modeling framework to understand how community composition impacts butyrate production in synthetic communities of prevalent and diverse human gut microbes. Butyrate production is a specialized function of a subset of species in the gut (∼10-25% of microbial genomes are predicted to harbor this pathway in healthy individuals^33^). By contrast, the production of other metabolic byproducts such as acetate and lactate are distributed more broadly across members of the gut microbiome. Thus, studying butyrate production allows us to investigate how ecological forces shape a function performed only by a specific subset of the community. Indeed, predictably modulating a specific function performed by a subset of organisms constitutes a core goal of microbiome engineering across different environments^34–36^.

To develop a system of microbes representing major metabolic functions in the gut, we selected 25 highly prevalent bacterial species from all major phyla in the human gut microbiome^37^. This community contained 5 butyrate producing Firmicutes which have been shown to play important roles in human health and protection from diseases (**Figure 2a, Table S1**). These 5 Firmicutes have the capability to ferment sugars and/or transform acetate to butyrate, allowing the recovery of NAD^+^ for further energy generation^38^. Additionally, *Anaerostipes caccae* (AC) can ferment lactate and acetate to butyrate, generating a modest amount of energy^39^. However, each of these species can alternatively produce acetate and/or lactate as fermentation products depending on the environmental context (**Figure 2b**).

**Figure 2.**
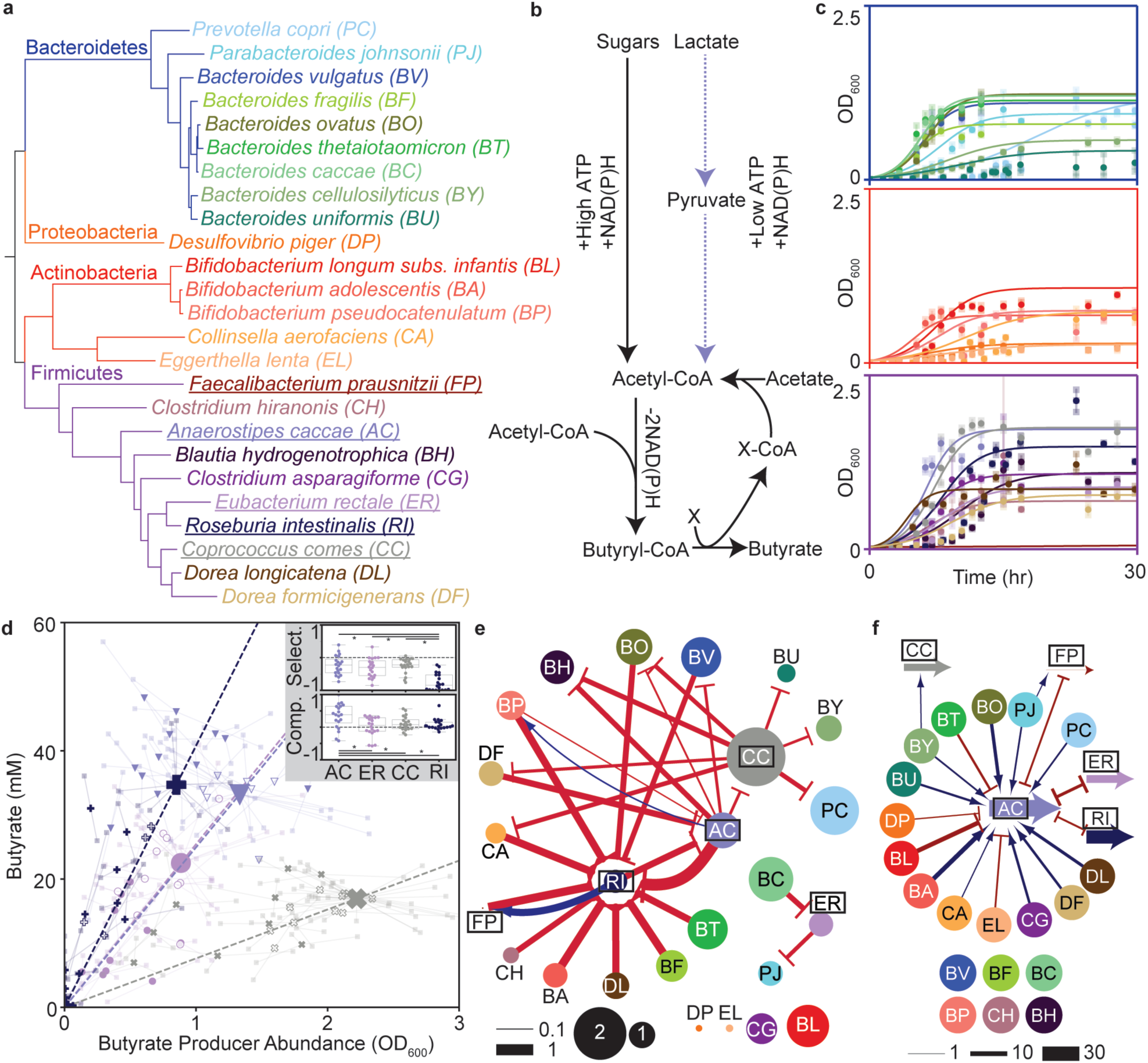
Characterizing interaction types in two-species communities. **(a)** Phylogenetic tree of the synthetic human gut microbiome community composed of 25 highly prevalent and diverse species. Branch color indicates phylum and underlined species denote butyrate producers. **(b)** Metabolic pathways for the transformation of sugars, acetate, and lactate into butyrate. Conversion of sugars or lactate to acetyl-CoA generates ATP and NAD(P)H, with higher ATP production per NAD(P)H from sugars. NAD(P)H is oxidized through conversion of acetyl-CoA to butyryl-CoA. Many substrates (X) can be used to exchange CoA between acetate and/or butyrate. In our system, *Anaerostipes caccae* has the unique capability to utilize the lactate conversion pathway (purple dashed arrows). **(c)** Monospecies growth responses over time. Transparent symbols indicate biological replicates connected to the corresponding mean (solid symbols) by transparent lines. Solid lines represent the generalized Lotka-Volterra (gLV) model fit to the data. Each plot shows the growth curves for species within the Bacteroidetes (top), Actinobacteria/Proteobacteria (middle) or Firmicutes (bottom) phylum. **(d)** Scatter plot of butyrate producer abundance and butyrate concentration for all pairwise communities containing at least one butyrate producer. Solid symbols indicate the mean of biological replicates of a community. Large symbols indicate butyrate producer monoculture. Smaller symbols indicate two-species communities, with closed symbols denoting significant differences in butyrate concentration and/or butyrate-producer abundance from the monoculture (p<0.05, t-test, unequal variance). Transparent squares indicate biological replicates and are connected to the corresponding mean with lines. Dashed lines indicate the predicted butyrate concentration assuming a constant butyrate yield based on monoculture data. Inset: distribution of selection and complementarity effects normalized by monoculture butyrate concentration for two-species communities. Asterisks indicate significant difference in the mean across butyrate producers (p<0.05, t-test, unequal variance) **(e)** Network representation of the inferred gLV inter-species interaction network based on data from **b** and **c**. Nodes size represents the abundance of each species in monoculture (OD_600_) at 48 hr and edges indicate interaction parameters with widths proportional to magnitude (units of hr^-1^ OD_600_^-1^) and color indicating sign (red negative, blue positive). Only edges with >95% confidence in sign are shown. **(f)** Network representation of regression model trained on data from **b** and **c**. Butyrate producer arrows denote monoculture butyrate production, nodes indicate non-butyrate producers, and edges represent modification of butyrate production in two-species communities. Edges connecting two butyrate producer arrows appear as bidirectional arrows since the directionality of the effect cannot be inferred. Edge widths are proportional to butyrate production (units of mM Butyrate). Only interactions with magnitude greater than 2 mM are shown.

Due to lack of defined media that universally support growth of gut microbes, most *in vitro* studies use rich media, making it difficult to interrogate the effects of unknown components on community behaviors^40^. To maximize our knowledge of the substrates available to the communities in our experiments and to simplify the metabolite quantification, we developed a single chemically defined medium that supports the growth of all species in monoculture with the exception of *Faecalibacterium prausnitzii* (FP) (**Methods**). We measured time-resolved growth of each species and constructed a gLV null model that assumed no inter-species interactions. Our results demonstrated a wide variety of growth dynamics within each phylum, including disparate growth rates and carrying capacities (**Figure 2c**). Using this system, we implemented an iterative design, test, learn (DTL) cycle (**Figure 1b**) to explore a vast community design space and explore an ecosystem functional landscape

### Butyrate production impacted by selection and complementarity

For the first cycle of our iterative DTL approach, we sought to decipher interactions impacting butyrate production in pairwise communities, with the goal of understanding how these interactions combine to shape community assembly and butyrate production in higher complexity communities. We grew each pairwise community containing at least one butyrate producer (the focal species of our system^41^) and measured species abundance and the concentrations of organic acid fermentation products (including butyrate, lactate, succinate and acetate) after 48 hours. Based on previous studies using pairwise communities to predict higher complexity community behaviors^6,14,42^, we hypothesized that these measurements would provide a highly informative dataset to develop an initial model that captured inter-species interactions shaping selection and complementarity effects in the system.

Based on our data, we first considered to what extent butyrate production was impacted by selection effects and complementarity effects using a model-free approach (**Figure 2d**). For each pairwise community, the selection effect was computed as the difference between the expected butyrate concentration assuming constant butyrate yield and the monoculture butyrate concentration (**Figure S1**). The complementarity effect was defined as the difference between the measured butyrate concentration of the community and the expected butyrate concentration assuming constant yield (**Figure S1, Methods**). Negative selection effects influenced all butyrate producers except FP, which did not grow in monoculture (**Figure 2d, inset**). Compared to the other butyrate producers, *Roseburia intestinalis* (RI) exhibited the largest negative selection effects, while AC tended to display positive complementarity effects. In sum, both selection and complementarity can modulate butyrate production, highlighting the utility of a building a composite model that captures both types of effects (**Figure 1a**). Further, the model-free approach to determining the contributions of selection and complementarity effects cannot be applied to communities containing multiple butyrate-producers. Therefore, our modeling approach can elucidate the selection and complementarity effects in communities with functional redundancy, representing real systems^33^.

To enable prediction of butyrate concentration in higher complexity communities, we used the data from the monoculture and pairwise community experiments to train our model (M1). The inferred gLV inter-species interaction network showed many negative interactions, including strong negative interactions impacting the growth of RI, consistent with the dominance of negative selection effects in our model-free analysis of RI pairwise communities (**Figure 2e**). A network representation of the parameters in the regression model indicated that AC had significantly more pairwise interaction terms than the other butyrate producers, consistent with the major role of positive complementarity effects in our model-free analysis of AC pairwise communities (**Figure 2f**).

### Model trained on pairwise consortia predicts 3-5 species community behaviors

To test our model’s ability to predict function in communities with an incremental increase in complexity, we implemented a second DTL cycle with the goal of mapping the functional landscape of 3-5 member communities. Model M1 was informed only by pairwise communities that contained at least one butyrate producer and was thus naïve to all interactions between non-butyrate producers. Therefore, we needed to make some assumptions to enable prediction of multi-species consortia containing combinations of non-butyrate producers. Based on patterns observed in previous gLV model parameter sets^6^, we hypothesized that unmeasured interactions could be estimated based on the trends in measured interactions across phylogenetic relatedness. Therefore, we used a matrix imputation method to estimate interaction parameters for unmeasured interactions in the gLV model (**Methods**). The resulting model was used to predict the probability distributions of butyrate production for all 3-5 species communities containing at least one butyrate producer (46,588 communities). The predicted butyrate production varied substantially between the combinations of butyrate-producer groups (**F**). To evaluate the ability of our model to predict the behaviors of butyrate producers in a variety of community contexts, we experimentally characterized 156 communities that spanned a broad range of predicted butyrate concentrations a (**Figure 3a**). The model prediction exhibited good agreement with the rank order of butyrate production (Spearman rho=0.84, p=9*10^−43^), though moderately overpredicted the magnitude on average (**Figure 3b**).

**Figure 3.**
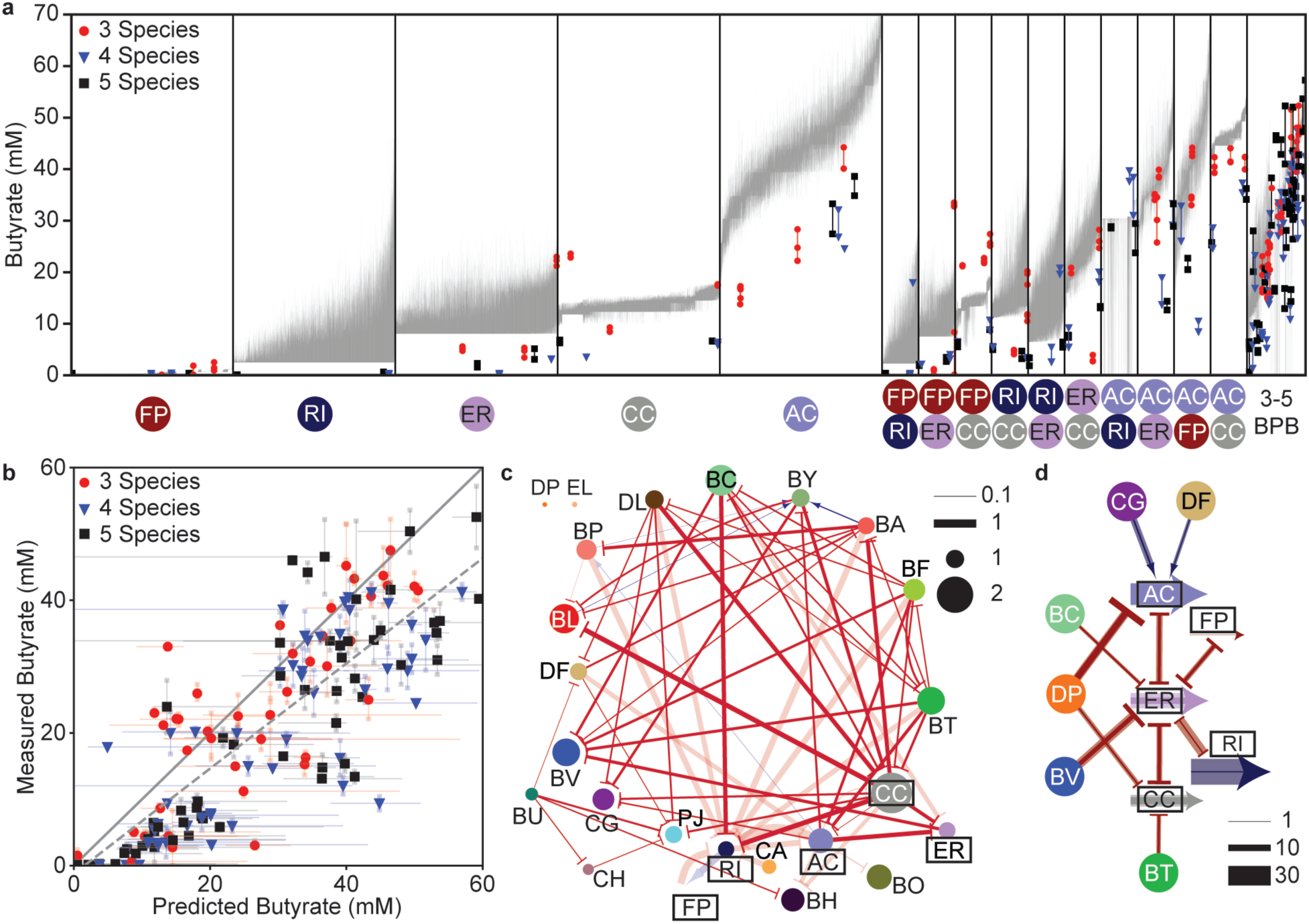
Model-guided investigation of low complexity synthetic human gut communities. **(a)** Predicted (grey bars) and measured (data points) butyrate concentrations for all 3-5 member communities containing at least one butyrate producer. Vertical black lines separate groups of communities based on the identities of the combination of butyrate producers specified on the x-axis. Communities with all combinations of 3-5 butyrate producers are included in the final bin for simplicity. Within each bin, communities are sorted in rank order of increasing median predicted butyrate production with a vertical grey bar indicating the 60 percent confidence interval of predictions for each community. Data points represent biological replicates of a selected subset of communities with replicates of a community connected by lines and the data point type indicating the number of species in each community (156 communities total). **(b)** Scatter plot of predicted and measured butyrate concentrations for communities in **a**. Transparent datapoints represent biological replicates of a community and are connected to the corresponding mean measurement (solid datapoints) by transparent lines. Prediction error bars indicate the 60 percent confidence interval of predicted butyrate. Solid grey line indicates x=y and dashed line indicates the linear regression between the mean measurement and median prediction (y=0.79x-1.2, Pearson r=0.83, p=6*10^−41^). Data point type indicates the number of species in each community. **(c)** Network representation of generalized Lotka-Volterra model updated with data from **b**. Node size represents the abundance of each species in monoculture (OD_600_), edge widths denote the magnitude of the inter-species interaction coefficients (units of hr^-1^ OD_600_^-1^) and color of the edges corresponds to the sign (red negative, blue positive). Faint interaction edges indicate interactions that did not change from the model trained on monospecies and pairwise communities (<2-fold change in magnitude of parameter mean). Only edges with >95% confidence in sign are shown. **(d)** Network representation of contributions of updated regression model to butyrate production in communities from **b**. Butyrate producer arrows indicate contribution to butyrate production independent of interactions. Nodes indicate non-butyrate producing species, and edges indicate modification of butyrate production in communities from **b**. Edges connecting two butyrate producers are bidirectional because it is not possible to discern which organism is producing the butyrate. For each butyrate producer, solid edge widths are proportional to the mean contribution and faint edge widths are proportional to the maximum contribution of the interaction across communities where those species were present (units of mM butyrate). Only interactions with maximum contribution >5 mM and with at least 4 communities including that interaction are shown.

The quality of these predictions demonstrated that our initial dataset was sufficient to build a model predicting broad trends in butyrate production but suggested that additional data was required to predict specific outliers. To understand the factors contributing to deviations between predicted and measured butyrate concentrations, we updated our models to decipher key inter-species interactions that model M1 failed to capture yielding model M2. The gLV model from M2 contained many new negative interaction parameters (46 new negative interactions, 15 conserved negative interactions) and sparse positive interactions (3 new and 2 conserved positive interactions) out of 386 possible observed interspecies interaction parameters in the designed set, primarily between non-butyrate producers (**Figure 3c**,). The regression model in M2 highlighted significant complementarity effects in the 3-5 member communities, with strong negative interactions between *Desulfovibrio piger* (DP)-AC and AC-*Eubacterium rectale* (ER) consistent with model M1 (**Figur 2**). We used the regression model with the experimental abundance measurements to quantify the magnitude and variability of each complementarity interaction across the experimentally measured communities due to differences in species growth. These data showed that some interactions consistently modified butyrate production in the presence of the species pair (e.g. DP-AC), whereas the contributions of other interactions to butyrate production varied across communities (e.g. ER-RI, *Clostridium asparagiforme* (CG)-AC) (**Figure 3d**). Equipped with this updated model, we set out to explore our model’s experimental design capabilities for communities of even greater complexity (i.e. >10 species).

### Model-guided exploration of complex community design space

One approach to determining the contributions of constituent community members to ecosystem behaviors involves characterization of the full and all single-species dropout consortia^6,26^. In our system, all 24 and the 25 member communities exhibited similar low butyrate production (∼10-15 mM Butyrate), except for a moderate increase of butyrate in the DP-lacking community (∼22 mM Butyrate) and a large decrease in the AC-lacking community (∼2 mM Butyrate) (**Figure S3**). Many of the 3-5 member communities displayed higher butyrate production than the highest complexity communities (**Figure 3b**), suggesting that high complexity communities may trend towards an undesired low butyrate producing state. Additionally, the concentrations of all measured organic acids spanned a much smaller range in the 24 and 25 member communities than the low complexity (1-5 member) assemblages (**Figure S3b**), further supporting the notion that communities of increasing complexity may trend toward a similar functional state. Indeed, a key challenge in engineering microbial communities is the tendency to assemble to compositional attractors and resist change, due to a multitude of abiotic and biotic interactions^43–46^. In our system, implementing a standard approach of analyzing the highest complexity set of consortia failed to elucidate diverse community metabolic states, highlighting the utility of the model to design sub-communities that span the functional space.

To address this challenge, we used our model M2 to design complex communities (>10 species) that deviated from the observed trend towards low butyrate production. Since the human gut microbiome exhibits functional redundancy in butyrate pathways^33^, we used model M2 to simulate the assembly of all communities containing all five butyrate producers to map species abundance to butyrate concentration (1,048,575 communities). Based on the hypothesis that high-complexity communities may trend towards low butyrate production, we found it useful to consider the full community as a reference frame, representing a potential compositional attractor state, when visualizing the relationship between community composition and butyrate production. Consistent with this notion, the model predicted the full community to have butyrate production similar to the average of all communities, with other communities diverging from this average behavior with increasing distance in composition (Euclidean distance between endpoint abundances) from the full community (**Figure 4a**). The landscape of communities was partitioned into two large clusters based on the presence or absence of the prevalent sulfate-reducing Proteobacteria DP^47^. Corroborating these results, DP had the strongest negative complementarity interaction with AC and CC in the regression model of M2 as well as a significant negative impact on butyrate production in the single-species dropout consortia (**Figure 3d, Figure S3a**). This inferred complementarity effect and predicted shift in the butyrate production landscape suggests that the presence of DP may substantially alter the metabolic activities shaping butyrate production.

**Figure 4.**
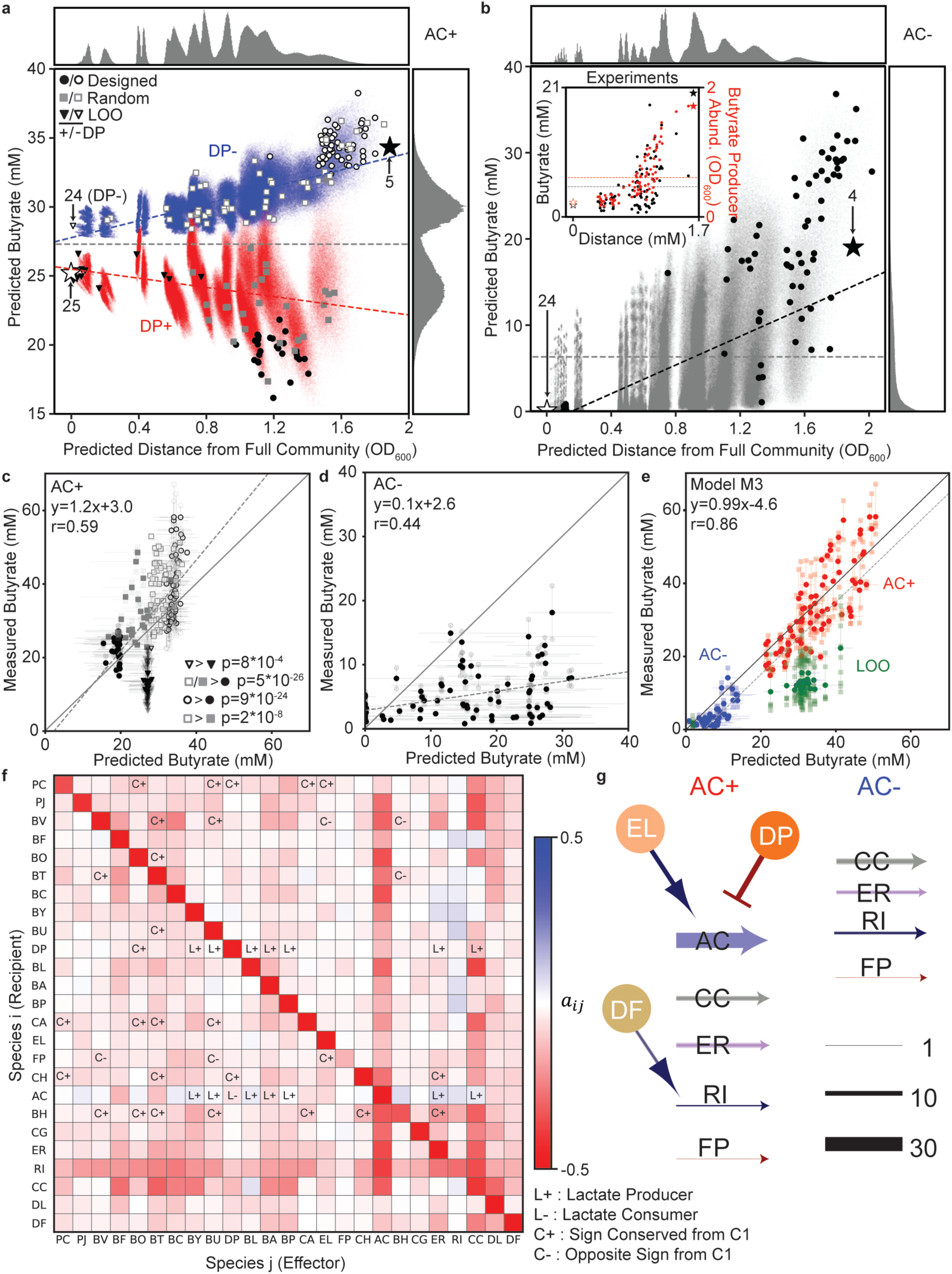
Model-guided exploration of butyrate production landscape. **(a)** Scatter plot of Euclidean distance in community absolute abundance from predicted full 25-member community versus predicted butyrate concentration for all possible communities. Histograms indicate the distribution of communities across the given axis. Communities are colored according to the presence (red) or absence (blue) of *Desulfovibrio piger* (DP). Blue and red dashed lines indicate the linear regression of communities with (red, y=-1.7x+25.5, r=-0.26) or without (blue, y=3.1x+27.8, r=0.72) DP. The white star indicates the full 25-member community and black star indicates the community of all butyrate producers. Large data points indicate communities chosen for experimental validation. Black triangles indicate leave-one-out communities, black circles indicate designed communities, and grey squares indicate random communities, with open/closed symbols indicating absence/presence of DP. **(b)** Scatter plot of Euclidean distance in community composition from predicted 24-member community excluding *Anaerostipes caccae* (AC) versus predicted butyrate concentration for all possible communities. Histograms indicate the distribution of communities across the given axis. Grey dashed line indicates the mean predicted butyrate concentration across all communities. Blue dashed line indicates the linear regression of all communities (y=8.3x-1.4, r=0.50). The white star indicates the full 24-member community and the black star indicates the 4 butyrate-producer community. Large data points indicate communities chosen for experimental validation. Inset: mean experimental measurements of butyrate concentration (black) and total abundance of butyrate producers (red) versus the distance from the full 24-member community. The grey and red dashed lines represent the mean butyrate concentration and total butyrate producer abundances across measured communities, respectively. **(c)** Scatter plot of predicted versus measured butyrate concentration for communities in **a**. Transparent data points indicate biological replicates and are connected to the corresponding mean values by transparent lines. Data points denote the median with error bars spanning the 60% confidence interval. Solid line indicates x=y. Dashed line indicates linear regression of median prediction versus mean measurement (y=1.2x+3.0, r=0.59, p=1*10^−18^). Legend indicates statistically significant differences in measured butyrate between populations of communities (Kruskal-Wallis test). **(d)** Scatter plot of predicted versus measured butyrate for communities in **b**. Transparent data points indicate biological replicates and are connected to the corresponding mean by transparent lines. Data points represent the median with error bars spanning the 60% confidence interval. Solid grey line indicates x=y. Dashed line indicates the linear regression of the median versus mean measurement (y=0.1x+2.6, r=0.44, p=2*10^−5^). **(e)** Scatter plot of predicted versus measured butyrate for complex communities using model M3. Transparent data points indicate biological replicates and are connected to the corresponding mean by transparent lines. Solid grey line indicates x=y. Dashed line indicates linear regression of prediction versus mean measurement (y=0.99x-4.6, r=0.86, p=1*10^−44^). **(f)** Heat-map of the median value of the inter-species interaction coefficients (a_ij_) for the M3 gLV model. Interactions impacting AC and DP are annotated with L+ or L-if species j produced or consumed >10 mM lactate in monoculture. Inter-species interactions included in the model community C1 from (Venturelli et al., Mol. Sys. Bio., 2018)^6^ are annotated with C+ or C-if interactions from both models had magnitudes greater than 0.05 hr^-1^ and had the same or opposite sign, respectively. **(g)** Network representation of updated M3 regression model. Butyrate producer arrows indicate contribution to butyrate production independent of inter-species interactions. Nodes indicate non-butyrate producing species, and edges indicate modification of butyrate production (blue, increased; red, decreased) in all complex communities (>10 species). Solid edge widths are proportional to the mean contribution and faint edge widths are proportional to the maximum contribution of the interaction across communities where those species were present (units of mM butyrate). Only interactions with maximum contribution >5 mM and with at least 4 communities including that interaction are shown.

We evaluated the capability of our model to guide broader exploration of the functional landscape and identify infrequent communities that deviate from the typical behavior by designing 28 low- and 54 high-butyrate communities each with 11-17 members and containing all 5 butyrate-producers. In addition, we randomly selected 82 communities with the same complexity constraints to evaluate whether our model-guided design procedure could elucidate a set of communities that spanned a broader range of metabolic states, scored by the variance in butyrate concentration (**Figure 4a**). The 82 designed communities exhibited a higher variance in mean butyrate production than the 82 random communities (designed communities s.d.=11 mM, random communities s.d.=8 mM, Levene test, p=0.043), demonstrating a major advantage of the model-guided approach for designing communities to broadly explore regions of the functional landscape (**F**). Consistent with our model predictions, communities containing DP exhibited lower butyrate compared to communities excluding DP in both the designed and the randomly chosen communities (**Figure 4c**). While the model predicted the rank order of butyrate concentrations in these communities moderately well (Spearman rho=0.67, p=3*10^−25^), some of the highest butyrate production communities were underpredicted by the model (**Figure 4c**).

### Selection effects dominate in high complexity communities lacking AC

In microbial consortia, the contributions of individual members to a given function can be broadly distributed, wherein key driver species can exhibit a substantially larger contribution to community-level functions than the other members^6,11,14^. In our system, the 24-member community lacking AC exhibited 1.9±1.0 mM butyrate (mean±s.d., n=8), substantially lower than any observed complex (>10 species) community containing AC and qualitatively consistent with our model which predicted no butyrate (**Figure 4b, Figure S3b**). Therefore, AC was a driver of butyrate production in complex communities. There are large interpersonal differences in gut microbiota composition due to environmental factors and host-microbe interactions^48^. Thus, some species such as AC may be not be present in certain individuals^49^. To evaluate the capability of our model to steer systems from low to high butyrate producing states independent of the presence of particular species, we designed high butyrate producing complex communities lacking the driver species AC.

To do so, we used model M2 to simulate all communities containing the four butyrate producers excluding AC (1,048,575 communities) to forecast species abundance and butyrate production. Similar to the 5 butyrate-producer case, we used the 24-member community lacking AC as a reference frame for quantifying deviations from a potential compositional attractor. While most communities were predicted to have low butyrate production, butyrate production increased with the distance from the 24-member community (**Figure 4b**). To explore this design space and evaluate whether our model could identify communities with low or high butyrate activity, we experimentally assembled 84 communities containing 11-19 members that were predicted to display a broad range of butyrate production capabilities (**Figure 4b**). Mirroring our model prediction, distance from the 24-member community in species composition was positively correlated with butyrate production (Spearman rho=0.56, p=3*10^−8^) as well as butyrate producer abundance (Spearman rho=0.85, p=1*10^−24^) (**Figure 4b**, inset). However, the model substantially overpredicted butyrate production (**Figure 4d**). Therefore, we sought to continue the DTL paradigm by training our model on the new data.

### Updated model predicts butyrate production in high complexity communities

The discrepancies between our model predictions and experimental measurements in complex communities were either due to missing information about certain pairwise interactions (i.e. poor parameter estimates due to unobservable interactions) or higher-order interactions that could not be captured by our pairwise model (i.e. model structure fails to represent system behaviors). To distinguish between these possibilities, we updated our model by training on a subset of high-complexity communities: the random set of communities containing all butyrate producers (82 communities) and a randomly sampled half of the communities lacking AC (42 communities) (**Fi**). Notably, the updated model M3 predicted the measurements of high-complexity communities with high accuracy, demonstrating that the pairwise model structures could explain the quantitative trends in the data when provided with sufficient information (**Figure 4e**). The predictive capability of the model required information from complex communities, supporting recent theoretical work suggesting that the typical pairwise community experimental design may not be the most efficient for building predictive models of complex systems^26^.

We next examined the changes in the inferred parameters between models M2 and M3 to provide insights into key microbial interactions impacting complex community behaviors. The major changes in the updated gLV M3 model were new values for all previously unobserved pairwise interactions as well as modification of previously observed interaction parameters (**Figure 4f, Figure S2f**). Negative interactions (<-0.05 hr^-1^ (OD_600_ Species j)^-1^) dominated the network, representing 49.8% of the interspecies interaction parameters. By contrast, only 1.7% of interactions were strong positive (>0.05 hr^-1^ (OD_600_ Species j)^-1^), consistent with previous observations of the prevalence of negative interactions in microbial communities^6,50^. Notably, 70% of the previously observed interspecies interaction parameters fell within the 60% confidence interval of the posterior distribution for model M2, demonstrating that our M2 model was accurate but lacked sufficient information to be highly confident in the estimated parameter values.

In the updated gLV model, species which secreted lactate in monoculture tended to have a positive impact on the growth of AC (**Figure 4f**). Although DP is also a lactate consumer^47^, it did not tend to benefit from monospecies lactate producers. This result highlights the benefits of using data from multiple levels of community complexity for training gLV models as these interactions were not captured by models M1 and M2, trained only on lower-order community contexts. To understand how inter-species interactions vary across chemical composition contexts, we compared the inferred inter-species interaction coefficients in the M3 gLV model to those from a previous study that used a gLV model to study a 12-member subset of our community (PC, BV, BO, BT, BU, DP, CA, EL, FP, CH, BH, and ER) in a different (rich) media^6^ and found that 27 parameters with magnitude >0.1 hr^-1^ (OD_600_ species j)^-1^ shared a sign and only 5 had opposite sign (**Figure 4f**). The high percentage (84%) of qualitatively consistent interaction coefficients inferred based on measurements in two different environmental contexts provides confidence in using parameterized gLV models as prior information to forecast system behaviors in new environments.

The regression model from M3 identified three interactions driving complementarity effects in the 5 butyrate producer communities including *Eggerthella lenta* (EL)-AC, DP-AC, and *Dorea formicigenerans* (DF)-RI (**Figure 4g**). In the absence of AC, substantial complementarity effects were not detected (**Figure 4g**), consistent with the absence of strong complementarity effects in lower-complexity communities lacking AC. In our system, AC has the unique capability to transform lactate to butyrate in addition to production of butyrate from sugars (**Figure 2b**), suggesting that metabolic flexibility may be a key determinant of complementarity effects. In sum, our modeling framework representing pairwise interactions could accurately predict community composition and butyrate production in complex communities and could be used to decipher key microbial interactions impacting metabolic outputs.

### Mechanistic insights identified from inferred interaction networks

We sought to analyze the patterns in our inferred interactions to identify mechanistic hypotheses about the potential ecological and molecular factors driving butyrate production. The low butyrate productivity of specific communities could stem from a global reduction in metabolic activities for the conversion of sugars to organic acid fermentation products. However, the amount of total carbon in acetate, lactate, and propionate was inversely proportional to the amount of carbon in butyrate in complex communities (**Figure S4**), indicating that metabolic tradeoffs dictated the production of specific organic acids. Therefore, we considered how interactions identified by our model could influence such tradeoffs.

We analyzed the inferred interaction networks to provide generalizable insights into metabolic processes impacting butyrate production in our system. We first considered the largest negative complementarity effect in our system between AC and DP (**Figure 4g**). While these two species have previously been shown to compete for lactate *in vitro*^51^, excess lactate was present in communities containing both DP and AC, suggesting that competition over limited lactate was not a major determinant of the negative complementarity effect (**Figure 5a**). In addition, a large negative complementarity effect was observed in the 3-5 member communities between DP and CC, which does not utilize lactate for butyrate production (**Figure 3d**).

**Figure 5.**
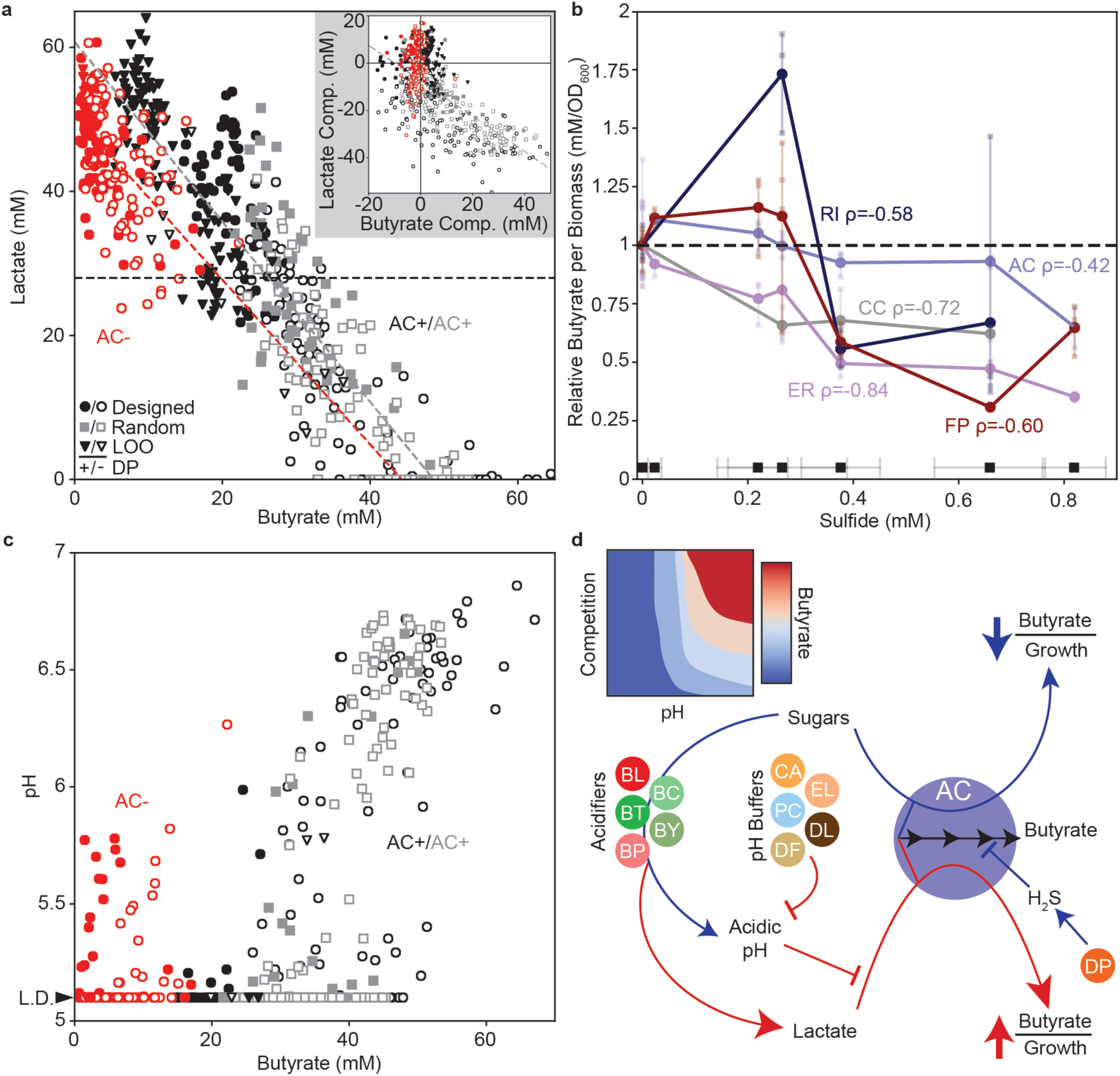
Model-guided identification of molecular mechanisms impacting butyrate production. **(a)** Scatter plot of butyrate concentration versus lactate concentration for complex communities (>10 species). Each data point indicates a biological replicate of a community. Grey dashed line indicates the linear regression for communities containing AC (y=-1.3x+61, r=-0.91, p=5*10^−182^), red dashed line indicates the linear regression for communities lacking AC (y=-1.1x+51, r=-0.56, p=8*10^−16^) and black horizontal dashed line indicates initial concentration of lactate in the media (28 mM). Inset: butyrate complementarity versus lactate complementarity. Grey dashed line indicates the linear regression for communities containing AC (y=-0.75x-7.5, r=-0.63, p=4*10^−53^). Pearson correlation for communities lacking AC was not statistically significant (p=0.12). **(b)** Butyrate concentration per unit biomass as a function of sulfide concentration. Butyrate yield per biomass was normalized to the no sulfide condition. Circles indicate the mean of biological replicates, with individual replicates shown as transparent squares. Black squares indicate the mean measured sulfide concentration for each treatment level with error bars indicating the standard deviation of at least 10 technical replicates. Species labels are accompanied by statistically significant Spearman correlation coefficients (ρ) between all biological replicates of that species and mean sulfide concentration for each level (p<0.05, AC p=0.02; CC p=0.002; ER p=3*10^−8^; RI p=0.02; FP p=0.0008). **(c)** Scatter plot of butyrate concentration versus pH for complex communities. Each data point indicates a biological replicate of a community. **(d)** Schematic representing proposed driving mechanisms impacting butyrate production by AC in complex communities. Red edges denote processes that negatively impact butyrate production and blue edges represent processes that enhance butyrate production. The abundance of species that acidify the environment were positively correlated with lactate concentration and negatively correlated with pH in complex communities. The abundance of pH buffering species were positively correlated with pH in complex communities. Note that species contributions to these processes are expected to be context-dependent. Inset: proposed qualitative butyrate landscape as a function of the strength of resource competition for sugars and environmental pH.

Since some *Desulfovibrio* species have the capability to use butyrate as an energy source^52^, we tested whether decreased butyrate in the presence of DP could be due to butyrate consumption. To investigate this hypothesis, we grew DP in media supplemented with different concentrations of sodium butyrate ranging between 0 and 100 mM and measured the butyrate concentration after 48 hours of incubation. The presence of DP did not alter the concentration of butyrate in any condition, suggesting that decreased butyrate due to consumption or degradation was not a major factor contributing to the negative complementarity effects associated with DP (**Figure S5**). One unique metabolic characteristic of DP in our system is the capability to reduce sulfate to hydrogen sulfide (H_2_S). Therefore, we hypothesized that H_2_S may contribute the negative impact of DP on butyrate production (**Figure 4a**). To test this hypothesis, each butyrate producer was grown in media supplemented with a range of sulfide concentrations. Notably, the butyrate production per unit biomass decreased with increasing sulfide concentration for all butyrate producers (**Figure 5b**). These data suggest that the levels of H_2_S produced by the host and constituent members of gut microbiota could shape butyrate production in the human gut microbiome.

We next investigated the factors that contribute to strong positive complementarity interactions influencing AC from EL or DF in complex communities with all butyrate producers (**Figure 4f**). Butyrate concentration exhibited a strong negative correlation with lactate concentration in complex communities (**Figure 5a**). Based on this correlation, we hypothesized that communities with higher butyrate concentration than expected based on monoculture butyrate yield (i.e. total positive butyrate complementarity) would exhibit lower lactate concentration than expected based on monoculture lactate yield (i.e. total negative lactate complementarity) (**Methods**). Our results demonstrated a negative correlation between butyrate and lactate complementarity in communities with AC, but not in communities without AC (**Figure 5a**, inset). These results suggest that the majority of excess butyrate that was not predicted based on monospecies butyrate yield was attributed to conversion of lactate to butyrate by AC, suggesting that this metabolic mode for butyrate production was driving the inferred complementarity effects. Thus, we next considered potential environmental factors that could inhibit the conversion of lactate to butyrate.

Previous studies have shown that the environmental pH has a major impact on organic acid production by gut microbiota^53–56^. For example, in batch cultures of fecal inocula, supplemented lactate was converted entirely to butyrate, propionate, and acetate at pH 5.9 and 6.4, but not at pH 5.2. This abrupt metabolic shift at low environmental pH was attributed to inhibition of lactate consumption by AC and *E. hallii* (a closely related lactate-consuming butyrate producer in the clostridial cluster XIVa)^55^. Consistent with these results, butyrate concentration and pH were positively correlated in complex communities with AC (Spearman rho=0.73, p=1*10^−57^) or without AC (Spearman rho=0.29, p=1*10^−4^), though the correlation was much stronger in communities with AC (**Figure 5c**). A positive correlation between butyrate and pH could be attributed to reduced acidification of the media on a per carbon basis because one butyrate molecule is produced from (or as an alternative to) two acetate molecules (**Figure 2b**). However, we postulate that in the presence of AC, a different mechanism drives the substantially stronger correlation between pH and butyrate, wherein high butyrate production was enabled by an environmental pH maintained above 5.9 b which lac n by AC was inhibited^55^) **Figure 5c**).

The abundance of EL and DF were both positively correlated with pH (**Figure S6**) and had positive complementarity effects in the regression model (**Figure 4g**), consistent with the potential role of pH in mediating positive complementarity effects. Further, EL had a unique environmental impact by increasing the pH in monoculture, suggesting that this mechanism could contribute to the inferred positive complementarity effect towards butyrate production (**Figure S6**). The environmental pH for monospecies did not forecast the correlations between species abundance and pH in complex communities. For example, *Dorea longicatena* (DL) and *Dorea formicigenerans* (DF) strongly acidify the media in monoculture but are positively correlated with pH in complex communities (**Figure S6**), highlighting a challenging problem in relating species composition to broad functions such as environmental pH modifications across community contexts.

In sum, we postulate that transformation of lactate into butyrate by AC was controlled by a combination of pH modification and resource competition (**Figure 5d**). Based on this mechanism, AC can switch between low and high butyrate producing states depending on the environmental pH and availability of sugars. In communities containing pH buffering species such as EL that maintain the pH above the threshold, the butyrate yield per biomass is dependent on the strength of competition for limited pools of sugars (high growth, low butyrate yield state). After sugars have been depleted, AC switches to a low growth and high butyrate yield metabolic state that transforms lactate into butyrate. The timing of the AC metabolic switch depends on the strength of resource competition in the community. In low pH environments, transformation of lactate to butyrate is inhibited and thus AC competes for limited sugars, resulting in butyrate production that is proportional to growth (i.e. no complementarity effects). Corroborating this notion, lactate-utilizing butyrate producers, including AC, have been shown to prefer glucose over lactate and produce ∼5x more butyrate per unit biomass when grown on lactate versus glucose^39^. Consistent with the proposed mechanism, the abundance of AC was negatively correlated with butyrate in conditions with an endpoint pH > 6 (**Figure S7**). Above this pH threshold, there exists a tradeoff between the biomass of AC and butyrate production depending on the proportion of AC biomass derived from sugars (i.e. high biomass, low butyrate) or lactate (i.e. low biomass, high butyrate) (**Figure 5d**). In sum, the proposed mechanism indicates that in a pH buffered community, resource competition over energy rich nutrients could enhance butyrate production by AC by triggering a shift in metabolism from a low to high butyrate producing state. Further, this hypothesis may explain why positive butyrate complementarity effects from pH-buffering species were not captured by the M1 and M2 models trained on lower-order communities as there were fewer species and thus a reduced strength of resource competition. This analysis highlights that an interpretable statistical model that maps community composition to function can provide key biological insights into ecological and molecular mechanisms driving community functions and illuminates key metabolic modes of ecological drivers of community functions.

## DISCUSSION

We demonstrated that community-level functions can be designed using a modeling framework that predicts community assembly (selection effects) and then maps community composition to function (complementarity effects). Our results showed that the capability for butyrate production can vary over a broad range (0-20 mM or 10-60 mM butyrate in the absence and presence of AC, respectively) by manipulating the presence/absence of diverse non-butyrate producing species, highlighting the critical role of microbial interactions in community-level functions. We used a DTL cycle to develop a predictive model of butyrate production by synthetic human gut microbiome communities which enabled the identification of key microbial interactions and insights into potential molecular mechanisms driving butyrate production. Our results demonstrated that accurate prediction of community function in complex multi-member consortia (i.e. >10 species) required measurements of communities at similar levels of complexity. Thus, the predictive capability of computational models of microbial communities could be improved by choosing communities that span the range of complexities of interest, rather than implementing a standard procedure of characterizing pairwise communities^6,14,42^. Consistent with this proposed experimental design approach, recent theoretical work has demonstrated a similar perspective^26^. While our approach lacks a host-interaction component, the mechanistic nature of insights derived from our model will enable future work to adapt our pipeline to predict community-level functions in the mammalian gut environment. For instance, DP has been previously associated with IBD^57^, attributed to its H_2_S activity inhibiting oxidation of short chain fatty acids by the host via short-chain acyl-CoA dehydrogenase^58^. However, an additional mechanism through which hydrogen sulfide producers could contribute to IBD is by inhibiting microbial production of the anti-inflammatory metabolite butyrate via the analogous bacterial short-chain acyl-CoA dehydrogenase. Indeed, a previous study demonstrated that cecal contents of gnotobiotic mice colonized with an 8-member community plus DP contained less propionate and elevated 3-hydroxybutyrate (upstream intermediate of butyrate production) compared to the 8-member community alone. In this study, the butyrate concentration did not vary between conditions, which could have been masked by host butyrate consumption as the concentration was very low for both communities (<1 mM)^47^. This could be explained by H_2_S inhibition of bacterial short chain acyl-CoA dehydrogenases in butyrate and propionate metabolic pathways, observed as accumulation of 3-hydroxybutyrate in the former case and decreased propionate production in the latter. Additionally, this mechanistic insight could explain associations between colitis and other sulfur-reducing bacteria, such as *Bilophila wadsworthia*^59^, which has been shown to be associated with reduced expression of microbial butyrate synthesis pathways in a mouse model of colitis^60^.

A major strategy for microbiome modulation involves administration of non-resident species predicted to perform a target beneficial function^61^, including butyrate-producing bacteria^32^. Due to the plasticity of microbial metabolism, our results demonstrate that it is important to consider both how the resident community will enable growth of supplemented butyrate-producing bacteria as well as promote the desired metabolic states. Indeed, our results showed that in the presence of AC, the abundance of functional strains may not correlate with community-level metabolic functions due to complementarity effects that modify microbial metabolic modes.

More broadly, our work provides a foundation for implementing model-guided procedures to design community properties and guide development of ecological and mechanistic hypotheses for a wide range of applications. Simple modifications can be made to this framework to accommodate different observed system behaviors. For instance, we modeled our system using two models incorporating only pairwise interaction terms. While this provided a high level of interpretability, it has a limited flexibility for studying higher-order interactions, which may play a critical role in shaping microbiome properties. Additionally, we focused on a predicting single function, whereas designing communities for multifunctionality may be desirable in many cases. Both of these limitations could be addressed by modifying our approach using alternative growth and function models, leveraging advances in machine learning^62^ or integrating information from genome-scale metabolic models^13^.

In this work, we constructed models of community dynamics and function in a single media. The gut microbiome is exposed to a wide range of dietary substrates and the temporal changes in resource availability can dramatically shape community composition^63^. Our approach could be adapted to represent the molecular environment as a design variable to allow simultaneous exploration of the community and chemical composition design spaces to better understand how the molecular environment shapes microbial community functions. In sum, our methods provide a flexible foundation to explore design strategies for building microbial communities with target functions from the bottom-up and to understand molecular and ecological mechanisms influencing community-level functions.

## METHODS

### Strain Maintenance and Culturing

All anaerobic culturing was carried out in an anaerobic chamber with an atmosphere of 2.5±0.5% H_2_, 15±1% CO_2_ and balance N_2_. All prepared media and materials were placed in the chamber at least overnight before use to equilibrate with the chamber atmosphere. The strains used in this work were obtained from the sources listed in **Table S2** and permanent stocks of each were stored in 25% glycerol at -80°C. Batches of single-use glycerol stocks were produced for each strain by first growing a culture from the permanent stock in anaerobic basal broth (ABB) media (HiMedia or Oxoid) to stationary phase, mixing the culture in an equal volume of 50% glycerol, and aliquoting 400 *μ*L into Matrix Tubes (ThermoFisher) for storage at -80°C. Quality control for each batch of single-use glycerol stocks included (1) plating a sample of the aliquoted mixture onto LB media (Sigma-Aldrich) for incubation at 37°C in ambient air to detect aerobic contaminants and (2) Illumina sequencing of 16S rDNA isolated from pellets of the aliquoted mixture to verify the identity of the organism. For each experiment, precultures of each species were prepared by thawing a single-use glycerol stock and combining the inoculation volume and media listed in **Table S2** to a total volume of 5 mL (multiple tubes inoculated if more preculture volume needed) for stationary incubation at 37°C for the preculture incubation time listed in **Table S2**. All experiments were performed in a chemically defined medium (DM38), the composition of which is provided in **Table S3**.

### Monoculture Dynamic Growth Quantification

Each species’ preculture was diluted to an OD_600_ of 0.0066 (Tecan F200 Plate Reader, 200 uL in 96-Well Microplate) in DM38 and aliquoted into 3 replicates of 1 mL each in a 96 Deep Well (96DW) plate and covered with a semi-permeable membrane (Diversified Biotech) for stationary incubation at 37°C. At each time point, samples were mixed and OD_600_ was measured by diluting an aliquot of each sample into phosphate-buffered saline (PBS) into the linear range of the plate reader.

### Community Culturing Experiments and Sample Collection

To produce all desired community combinations, each species’ preculture was diluted to an OD_600_ of 0.0066 in DM38. Community combinations were arrayed in 96DW plates by pipetting equal volumes of each species’ diluted preculture into the appropriate wells using a Tecan Evo Liquid Handling Robot inside an anaerobic chamber. Each 96DW plate was covered with a semi-permeable membrane and incubated at 37°C. After 48 hours, 96DW plates were removed from the incubator and samples were mixed. Cell density was measured by pipetting 200 *μ*L of each sample into one microplate and diluting 20 *μ*L of each sample into 180 *μ*L of PBS in another microplate and measuring the OD_600_ of both plates (Tecan F200 Plate Reader). We selected the value that was within the linear range of the instrument for each sample. 200 uL of each sample was transferred to a new 96DW plate and pelleted by centrifugation at 2400xg for 10 minutes. A supernatant volume of 180 *μ*L was removed from each sample and transferred to a 96-well microplate for storage at -20°C and subsequent metabolite quantification by high performance liquid chromatography (HPLC). Cell pellets were stored at -80°C for subsequent genomic DNA extraction and 16S rDNA library preparation for Illumina sequencing. In some experiments, 20 *μ*L of each supernatant was used to quantify pH using a phenol Red assay^64^. Phenol red solution was diluted to 0.005% weight per volume in 0.9% w/v NaCl. Bacterial supernatant (20 *μ*L) was added to 180 *μ*L of phenol red solution, and absorbance was measured at 560 nm (Tecan Spark Plate Reader). A standard curve was produced by fitting the Henderson-Hasselbach equation to fresh media with a pH ranging between 3 to 11 measured using a standard electro-chemical pH probe (Mettler-Toledo). We used the following equation to map the pH values to the absorbance measurements.

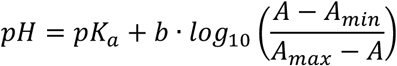

The parameters *b* and *pK*_*a*_ were determined using a linear regression between pH and the log term for the standards in the linear range of absorbance (pH between 5.2 and 11) with *A*_*max*_ representing the absorbance of the pH 11 standard, *A*_*min*_ denoting the absorbance of the pH 3 standard and *A* representing the absorbance of each condition.

### Sulfide Titration Experiment

Each species’ preculture was diluted to an OD_600_ of 0.0066 in DM38. FP cultures were supplemented with 1 g/L bacto yeast extract (BD) and 33 mM sodium acetate (Sigma Aldrich). Different volumes of a concentrated solution of sodium sulfide (Alfa Aesar) were added to the cultures to achieve the desired concentration and the cultures were incubated in capped 1.6 mL microfuge tubes for 24 hours at which point the OD_600_ was measured (Tecan F200 Plate Reader, 200 uL in 96-Well Microplate) and supernatants were collected for organic acid quantification via HPLC. Sulfide concentrations in the initial cultures were measured via the Cline assay^65^ to account for degradation of the sulfide stock during experimental setup. Briefly, 14.8 uL of Cline reagent was added to 185.2 uL of culture supernatant and incubated in a sealed 96-Well Microplate for 2 hours before diluting in 1% zinc acetate (Fisher) to the linear range of absorbance measurement at 667 nm (Tecan Spark Plate Reader). A standard curve was prepared similarly using sodium sulfide fixed in 1% zinc acetate. Cline reagent was prepared by dissolving 1.6 g N,N-dimethyl-p-phenylenediamine sulfate (Acros Organics) and 2.4 g FeCl_3_ (Fisher) in 100 mL 50% v/v HCl (Fisher) in water.

### HPLC Quantification of Organic Acids

Supernatant samples were thawed in a room temperature water bath before addition of 2 *μ*L of H_2_SO_4_ to precipitate any components that might be incompatible with the running buffer. The samples were then centrifuged at 2400xg for 10 minutes and then 150 *μ*L of each sample was filtered through a 0.2 *μ*m filter using a vacuum manifold before transferring 70 *μ*L of each sample to an HPLC vial. HPLC analysis was performed using either a ThermoFisher (Waltham, MA) Ultimate 3000 UHPLC system equipped with a UV detector (210 nm) or a Shimadzu HPLC system equipped with a SPD-20AV UV detector (210 nm). Compounds were separated on a 250 x 4.6 mm Rezex© ROA-Organic acid LC column (Phenomenex Torrance, CA) run with a flow rate of 0.2 ml min^-1^ and at a column temperature of 50°C. The samples were held at 4°C prior to injection. Separation was isocratic with a mobile phase of HPLC grade water acidified with 0.015 N H_2_SO_4_ (415 µL L^-1^). At least two standard sets were run along with each sample set. Standards were 100, 20, and 4 mM concentrations of butyrate, succinate, lactate, and acetate, respectively. For most runs, the injection volume for both sample and standard was 25 µl. The resultant data was analyzed using the Thermofisher Chromeleon 7 software package.

### Genomic DNA Extraction and Sequencing Library Preparation

Genomic DNA was extracted from cell pellets using a modified version of the Qiagen DNeasy Blood and Tissue Kit protocol. First, pellets in 96DW plates were removed from -80°C and thawed in a room temperature water bath. Each pellet was resuspended by pipette in 180 *μ*L of enzymatic lysis buffer (20 mM Tris-HCl (Invitrogen), 2 mM Sodium EDTA (Sigma-Aldrich), 1.2% Triton X-100 (Sigma-Aldrich), 20 mg/mL Lysozyme from chicken egg white (Sigma-Aldrich)). Plates were then covered with a foil seal and incubated at 37°C for 30 minutes with orbital shaking at 600 RPM. Then, 25 *μ*L of 20 mg mL^-1^ Proteinase K (VWR) and 200 *μ*L of Buffer AL (QIAGEN) were added to each sample before mixing with a pipette. Plates were then covered by a foil seal and incubated at 56°C for 30 minutes with orbital shaking at 600 RPM. Next, 200 *μ*L of 100% ethanol (Koptec) was added to each sample before mixing and samples were transferred to a Nucleic Acid Binding (NAB) plate (Pall) on a vacuum manifold with a 96DW collection plate. Each well in the NAB plate was then washed once with 500 uL Buffer AW1 (QIAGEN) and once with 500 *μ*L of Buffer AW2 (QIAGEN). A vacuum was applied to the Pall NAB plate for an additional 10 minutes to remove any excess ethanol. Samples were then eluted into a clean 96DW plate from each well using 110 *μ*L of Buffer AE (QIAGEN) preheated to 56°C. Genomic DNA samples were stored at -20°C until further processing.

Genomic DNA concentrations were measured using a SYBR Green fluorescence assay and then normalized to a concentration of 1 ng *μ*L^-1^ by diluting in molecular grade water using a Tecan Evo Liquid Handling Robot. First, genomic DNA samples were removed from -20°C and thawed in a room temperature water bath. Then, 1 *μ*L of each sample was combined with 95 *μ*L of SYBR Green (Invitrogen) diluted by a factor of 100 in TE Buffer (Integrated DNA Technologies) in a black 384-well microplate. This process was repeated with two replicates of each DNA standard with concentrations of 0, 0.5, 1, 2, 4, and 6 ng *μ*L^-1^. Each sample was then measured for fluorescence with an excitation/emission of 485/535 nm using a Tecan Spark plate reader. Concentrations of each sample were calculated using the standard curve and a custom Python script was used to compute the dilution factors and write a worklist for the Tecan Evo Liquid Handling Robot to normalize each sample to 1 ng *μ*L^-1^ in molecular grade water. Samples with DNA concentration less than 1 ng *μ*L^-1^ were not diluted. Diluted genomic DNA samples were stored at -20°C until further processing.

Amplicon libraries were generated from diluted genomic DNA samples by PCR amplification of the V3-V4 of the 16S rRNA gene using custom dual-indexed primers (**Table S3**) for multiplexed next generation amplicon sequencing on Illumina platforms (Method adapted from Venturelli et al. *Mol. Sys. Bio*., 2018). Primers were arrayed in skirted 96 well PCR plates (VWR) using an acoustic liquid handling robot (Labcyte Echo 550) such that each well received a different combination of one forward and one reverse primer (0.1 *μ*L of each). After liquid evaporated, dry primers were stored at -20°C. Primers were resuspended in 15 *μ*L PCR master mix (0.2 *μ*L Phusion High Fidelity DNA Polymerase (Thermo Scientific), 0.4 *μ*L 10 mM dNTP Solution (New England Biolabs), 4 *μ*L 5x Phusion HF Buffer (Thermo Scientific), 4 *μ*L 5M Betaine (Sigma-Aldrich), 6.4 *μ*L Water) and 5 *μ*L of normalized genomic DNA to give a final concentration of 0.05 *μ*M of each primer. Primer plates were sealed with Microplate B seals (Bio-Rad) and PCR was performed using a Bio-Rad C1000 Thermal Cycler with the following program: initial denaturation at 98°C (30 s); 25 cycles of denaturation at 98°C (10 s), annealing at 60°C (30 s), extension at 72°C (60 s); and final extension at 72°C (10 minutes). 2 *μ*L of PCR products from each well were pooled and purified using the DNA Clean & Concentrator (Zymo) and eluted in water. The resulting libraries were sequenced on an Illumina MiSeq using a MiSeq Reagent Kit v3 (600-cycle) to generate 2×300 paired end reads.

### Bioinformatic Analysis for Quantification of Species Abundance

Sequencing data were demultiplexed using Basespace Sequencing Hub’s FastQ Generation program. Custom python scripts were used for further data processing (Method adapted from Venturelli et al. Mol. Sys. Bio., 2018)^6^. Paired end reads were merged using PEAR (v0.9.10)^66^ after which reads without forward and reverse annealing regions were filtered out. A reference database of the V3-V5 16S rRNA gene sequences was created using consensus sequences from next-generation sequencing data or Sanger sequencing data of monospecies cultures. Sequences were mapped to the reference database using the mothur (v1.40.5)^67^ command classify.seqs (Wang method with a bootstrap cutoff value of 60). Relative abundance was calculated as the read count mapped to each species divided by the total number of reads for each condition. Absolute abundance of each species was calculated by multiplying the relative abundance by the OD_600_ measurement for each sample. Samples were excluded from further analysis if they had OD_600_>0.1 and they had less than 1000 total reads or >1% of the reads were assigned to a species not expected to be in the community.

### Model-Free Quantification of Complementarity

We quantified the contribution of complementarity effects to butyrate and lactate production in each community by calculating the difference between the measured metabolite concentration and the expected metabolite concentration based on monoculture yield according to the following equation:

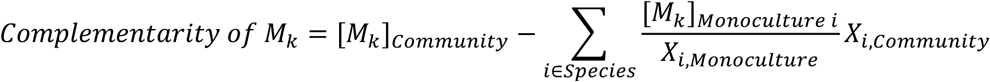

The variables *M*_*k*_ represents metabolite *k* (e.g. butyrate or lactate), [*M*_*k*_]_*community*_ represents the concentration of metabolite *k* measured in the community, [*M*_*k*_]_*monoculture i*_ denotes the concentration of metabolite *k* in the monoculture of species *i, X*_*i monoculture*_ represents the absolute abundance of species *i* in monoculture, *X*_*i community*_ is the absolute abundance of species *i* in the community, and the summation is across all species in the community.

### gLV Models and Training

We used a model with two modules: the gLV model to predict composition of the assembled community and a regression model with interaction terms to predict butyrate production as a function of the predicted community composition (**Figure 1a**). The gLV model is a set of *N* coupled first-order ordinary differential equations, where *N* denotes the number of species, of the form:

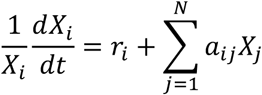

The species *X*_*i*_ is the abundance of species *i, r*_*i*_ is a parameter that represents the basal growth rate of species *i*, and *a*_*ij*_ is a parameter that represents interactions by modifying the growth rate of species *i* proportional to the abundance of species *j*. To prevent unbounded growth, *a*_*ij*_ is constrained to be negative when *i=j*, representing intra-species competition. This model has previously been used to understand and predict the behavior of complex microbial communities^6^ and provides an interpretable model form (e.g. which interspecies interactions are important) without introducing an excessive number of parameters (e.g. complex mechanistic models^68^).

We used a Bayesian parameter inference approach to estimate parameters for the gLV model from experimental measurements (adapted from Shin et al., *PLoS Computational Biology*, 2019^30^). Briefly, our method has a prior distribution for each model parameter and then varies the parameters to fit the model to the measured species abundances (mean of biological replicates) while penalizing deviations from the parameter prior distributions. These penalties provide a regularization effect, which is necessary when the model is underdetermined. We used L2 regularization because we expected inter-species competition to be prevalent and thus did not expect many interaction parameters to be negligible. After an optimal parameter set is found, this process is repeated hundreds of times after applying random noise to the experimental data proportional to the measured experimental variance to generate an ensemble of parameter sets (i.e. the posterior distribution). This posterior distribution is then used as the prior distribution when updating the model with new data. We adapted a previous implementation of this method in Julia for this work.

Before training the model on any data, we assumed a normally distributed prior for each parameter with mean of 0 and standard deviation equal to 1. We then trained the gLV model on time-series measurements of monoculture growth for each species, estimating a posterior distribution for each *r*_*i*_ and *a*_*ii*_ parameter (other *a*_*ij*_ posterior distributions were equal to the prior distribution). We used this posterior distribution as a prior distribution to update the model with the pairwise community data and generated the gLV module of Model M1, where posterior distributions were estimated from experimental data for *r*_*i*_, *a*_*ii*_, and *a*_*ij*_ where species *i* and species *j* co-occurred in the experimental data and the posterior distribution of *a*_*ij*_ for unobserved pairs was equal to the prior. We similarly updated the model using the 3 to 5-member community experiments to generate Model M2. Regularization coefficients for each iteration of the model updating process are shown in **Table S4**.

The gLV modules of Models M1 and M2 were underdetermined due to pairs of species never being observed in the same community within the training dataset. To generate parameters for these unobserved interactions, we used a matrix imputation approach to estimate the interaction parameters informed by the phylogenetic relatedness of species. First, we sorted the *a*_*i*j_ interaction parameter matrix such that the rows and columns occurred in the same order as the phylogenetic tree (**Figure 2a**). Next, we used K-nearest neighbors matrix imputation with K = 2 to estimate interaction parameters for species that were not observed in the training data (implemented in Python 3 using the fancyimpute package, https://pypi.org/project/fancyimpute/). This process was repeated independently for each parameter set in the posterior distribution.

While the parameter optimization portion of this model-training process had previously been found to scale with increasing number of pairwise community datasets^30^, we found that the optimization problem became intractable when attempting to estimate parameters from complex community data (i.e. >10 species). To address this problem, we used the nonlinear programming solver FMINCON in MATLAB to generate the gLV module of Model M3 by training on all data simultaneously. Using this method, the cost function for the optimization algorithm is computed using an ODE solver to simulate each community and the sum of mean squared errors for the community is computed and added to a L2 regularization term penalizing the magnitude of the parameter vector. To ensure that the model did not sacrifice the goodness of fit to the time-series monospecies data, the mean squared errors for these data were weighted more highly. The resulting optimization function was as follows:

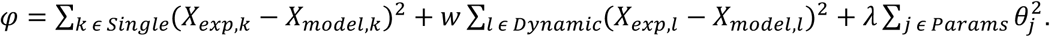

In this equation, single denotes the set of experiments where only the end point community composition was measured, dynamic indicates the set of time-series monospecies measurements, *w* is the weighting factor the time-series monospecies measurements, and *λ* represents the regularization coefficient. The FMINCON function identifies a parameter estimate which minimizes the cost function. We provided the median parameter values from Model M2 as an initial guess for the FMINCON function. We repeated this process with various values of *λ* and *w* to find a parameter set that simultaneously fits the Dynamic and Single datasets with maximal regularization penalty to prevent overfitting to the data (**Table S4**). We used a procedure based on the one described above for the Julia implementation to generate an ensemble of parameter sets (i.e. the posterior distribution) using FMINCON. Because each iteration of the FMINCON parameter estimation took several hours to complete, we massively parallelized the generation of each of the hundreds of parameter sets in the ensemble using resources from the UW-Madison Center for High Throughput Computing.

### Regression Models and Training

We used a regression model to represent a microbial community function with interaction terms:

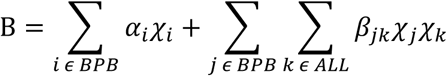

The variable B is the predicted butyrate concentration, *α*_*i*_ are parameters corresponding to each of the variables *χ*_*i*_ (end point abundances and time 0 presence or absence (1 or 0) for each butyrate producer, 10 variables total), and *β*_*jk*_ are interaction parameters corresponding to each pair of variables *χ*_*j*_ (end point abundances and time 0 presence or absence (1 or 0) for each butyrate producer,10 variables total) and *χ*_*k*_ (end point abundances and time 0 presence or absence (1 or 0) for all species, 50 variables total), excluding cases where *χ*_*j*_ and *χ*_*k*_ refer to the same species (450 total parameters). Model fitting was performed using custom scripts written in MATLAB and Python. We used L1 regularization to minimize the number of nonzero parameters. Regularization coefficients were chosen by using 10-fold cross validation and choosing the coefficient value with the lowest median mean-squared error for the test data. For models M1 and M2, ensembles of regression models were generated, one for each possible combination of butyrate producers, where samples containing butyrate producers from outside of each set were excluded. In this case, butyrate production from less productive species (e.g. FP) were small compared to more productive species (e.g. AC, ER, RI, CC) thus reducing the model accuracy for communities lacking the high productivity species. For Model M3, one regression model was generated using all data because all communities of interest contained highly productive butyrate producers.

### Model Simulations to Predict New Communities

Custom MATLAB scripts were used to predict community assembly and butyrate production, for many communities as described in the text (e.g. all communities containing all 5 butyrate producers for **Figure 4a**). For each community, the growth dynamics were simulated using each parameter set from the posterior distribution of the gLV model. The resulting community compositions for each simulation were an input to the regression model to predict butyrate concentration. Statistics on the resulting distributions of butyrate concentration and abundance of each species were stored for later plotting. Because of the large number of communities and the large number of parameter sets (i.e. hundreds of simulations per community), we used parallel computing (MATLAB parfor) to complete the simulations in a reasonable timeframe (∼4 days for the communities in **Figure 4a**).

## ACKNOWLEDGEMENTS

We would like to thank Sungho Shin, Jordan Jalving, and Victor Zavala for their advice related to implementing Julia parameter estimation methods. In addition, we are grateful to Mayank Baranwal and Alfred Hero for conversations which inspired the matrix imputation approach for estimating unobserved interaction parameters. We would like to thank Federico Rey for generously taking the time to provide advice that improved the manuscript. Research was sponsored by the National Institutes of Health and was accomplished under Grant Number R35GM124774 and University of Wisconsin-Madison Office of the Chancellor and Vice Chancellor for Research and Graduate Education with funding from the Wisconsin Alumni Research Foundation. S.E.H. was supported by the National Institute of General Medical Sciences of the National Institutes of Health under Award Number T32GM008349. R.L.C. was supported in part by an NHGRI training grant to the Genomic Sciences Training Program (T32 HG002760). This research was performed using the computing resources and assistance of the UW-Madison Center for High Throughput Computing (CHTC) in the Department of Computer Sciences. The CHTC is supported by UW-Madison, the Advanced Computing Initiative, the Wisconsin Alumni Research Foundation, the Wisconsin Institutes for Discovery, and the National Science Foundation, and is an active member of the Open Science Grid, which is supported by the National Science Foundation and the U.S. Department of Energy’s Office of Science.

## AUTHOR CONTRIBUTIONS

O.S.V. and R.L.C conceived the study. R.L.C., J.J.H., S.E.H., and B.M.C. carried out the experiments. R.L.C. implemented computational modeling. R.L.C., S.E.H. and O.S.V. analyzed the data. B.M.C. proposed inhibition of butyrate production by hydrogen sulfide. D.A.N. and D.M.S. designed and implemented metabolite measurements. O.S.V. secured funding. R.L.C. and O.S.V. wrote the paper and all authors provided feedback on the manuscript.

