## Supplementary Information for "Design of synthetic human gut microbiome assembly and function"

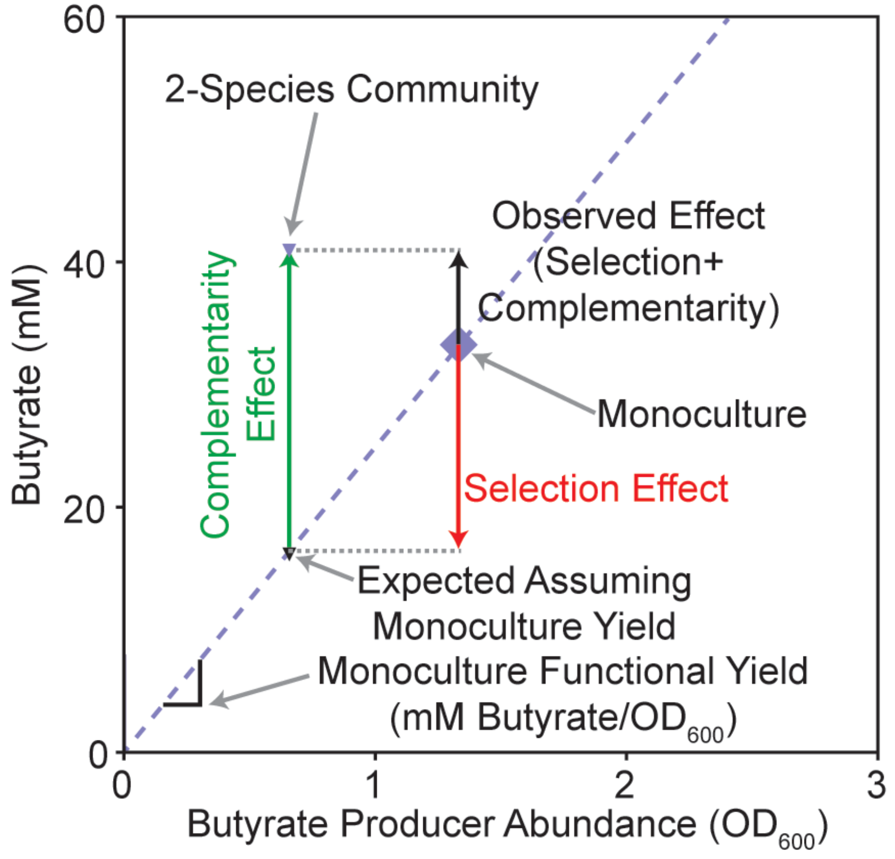


**Figure S1. Model-free analysis of selection and complementarity effects in pairwise consortia.** Representative comparison between a pairwise community (small purple triangle) to the corresponding butyrate producer monoculture (large purple diamond). Dashed purple line indicates expected concentration of butyrate as a function of butyrate producer abundance assuming monoculture functional yield. Small black triangle falls on this line at the butyrate producer abundance in the pairwise community. The observed change in butyrate concentration in the pairwise community (difference between pairwise community butyrate and monoculture butyrate, black arrow) can be decomposed into the sum of the selection effect (difference between monoculture butyrate and expected butyrate assuming monoculture yield, red arrow) and the complementarity effect (difference between pairwise community butyrate and expected butyrate assuming monoculture yield, green arrow).

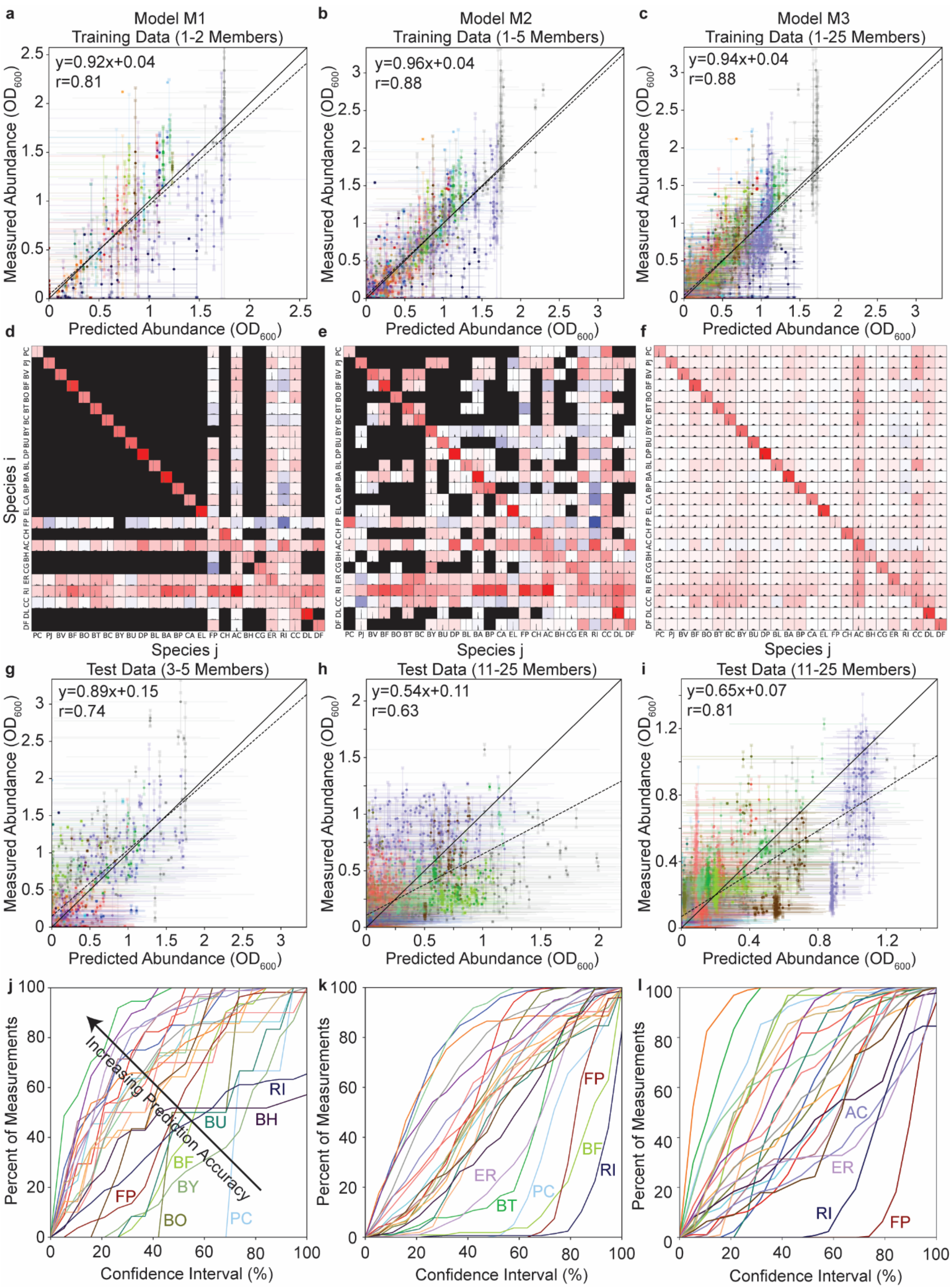
a
Model M1
Training Data (1-2 Members)
y=0.92x+0.04
r=0.81
Measured Abundance (OD600)
Predicted Abundance (OD600)
b
Model M2
Training Data (1-5 Members)
y=0.96x+0.04
r=0.88
c
Model M3
Training Data (1-25 Members)
y=0.94x+0.04
r=0.88
d
e
f
Species i
Species j
PC PJ BV BF BO BT BC BY BU DP BL BA BP CA EL FP CH AC BH CG ER RI CC DL DF
g
Test Data (3-5 Members)
y=0.89x+0.15
r=0.74
h
Test Data (11-25 Members)
y=0.54x+0.11
r=0.63
i
Test Data (11-25 Members)
y=0.65x+0.07
r=0.81
j
Increasing Prediction Accuracy
RI
BH
BU
BF
BY
FP
BO
PC
Percent of Measurements
Confidence Interval (%)
k
FP
BF
RI
ER
BT
PC
l
AC
ER
RI
FP

**Figure S2. Prediction of community dynamics using the Lotka-Volterra (gLV) model. (a-c)** Scatter plots of predicted versus measured species abundance for the training data for models M1-M3. Solid data points indicate the mean measured abundance and median predicted abundance for each species in each community. Transparent data points indicate biological replicate measurements and are connected to the corresponding mean values with lines. For **a** and **b**, horizontal error bars indicate the 60% confidence interval of the model prediction distribution. Colors correspond to legend in **Figure 2a.** Dashed line indicates the linear regression between the mean measured abundances and the median predicted abundances. **(d-f)** Heat-maps of model parameters $a_{ij}$, which quantifies the impact of species *j* on the growth rate of species *i*. Color of the square indicates the median parameter value (red is negative, blue is positive). Histograms within each subplot indicate the distribution of parameter values from the inference analysis. Solid black subplots indicate pairs that were not present in the corresponding training data. **(g-i)** Scatter plots of predicted versus measured species abundance for the test data (i.e. communities not included in the training set). **(j-l)** Percentage of measurements of the abundance of each species (colors correspond to legend in **Figure 2a**) that fall within a given confidence interval. Arrows indicate the direction of increasing model prediction accuracy.

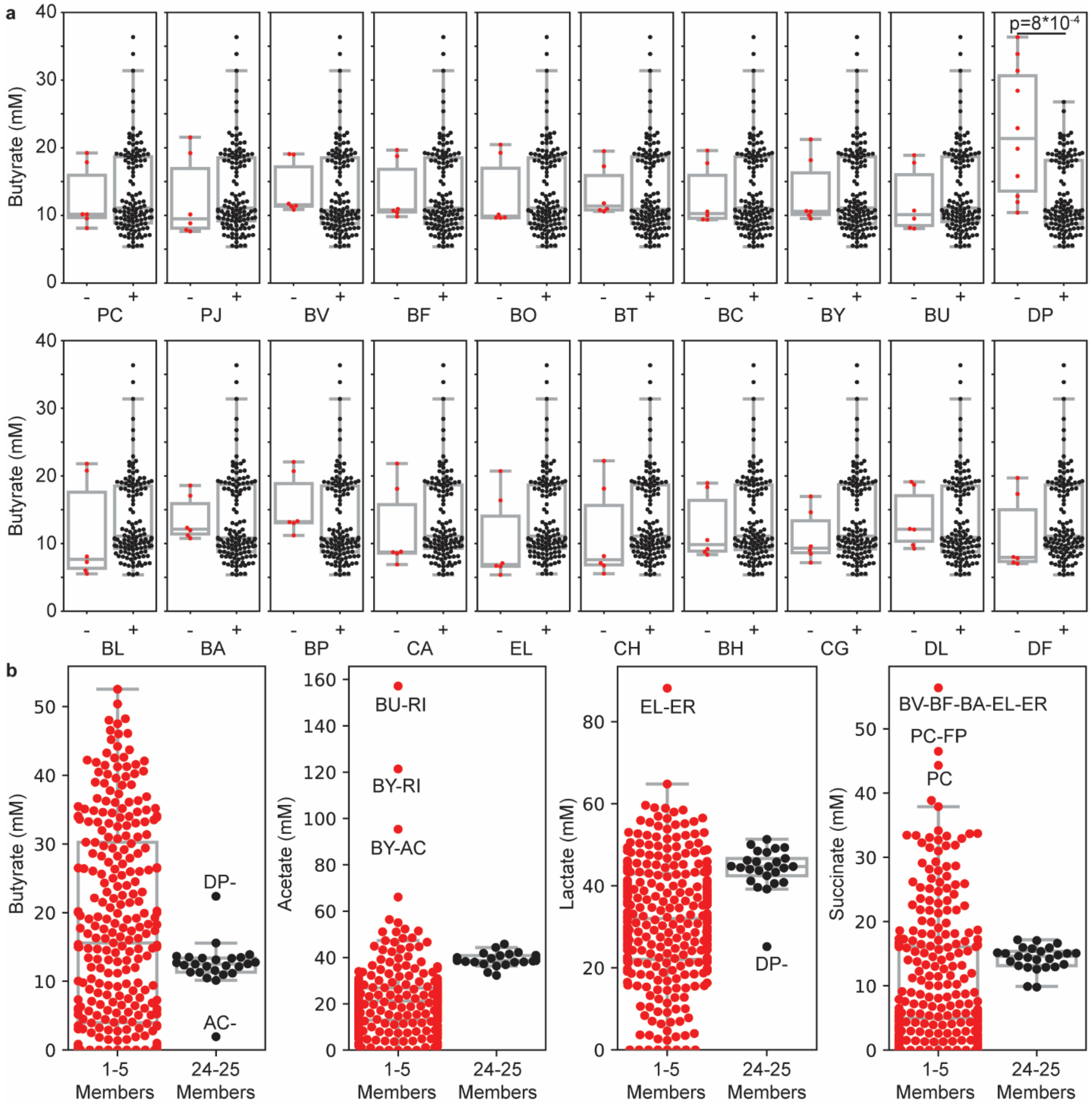


**Figure S3. Metabolite concentrations at different levels of community complexity. (a)** Butyrate production in single-species dropout communities (24-25 members). Each subplot shows a comparison of all 24-25 member communities that include all 5 butyrate producers with (+) or without (-) the indicated species. Each data point indicates a biological replicate of a community. *Desulfovibrio piger* (DP) is the only species with a statistically significant difference ($p<0.001$). **(b)** Distribution of organic acid concentrations in low complexity communities (1-5 members) versus single-species dropout consortia (24-25 members). Each datapoint indicates the mean organic acid concentration for each community.

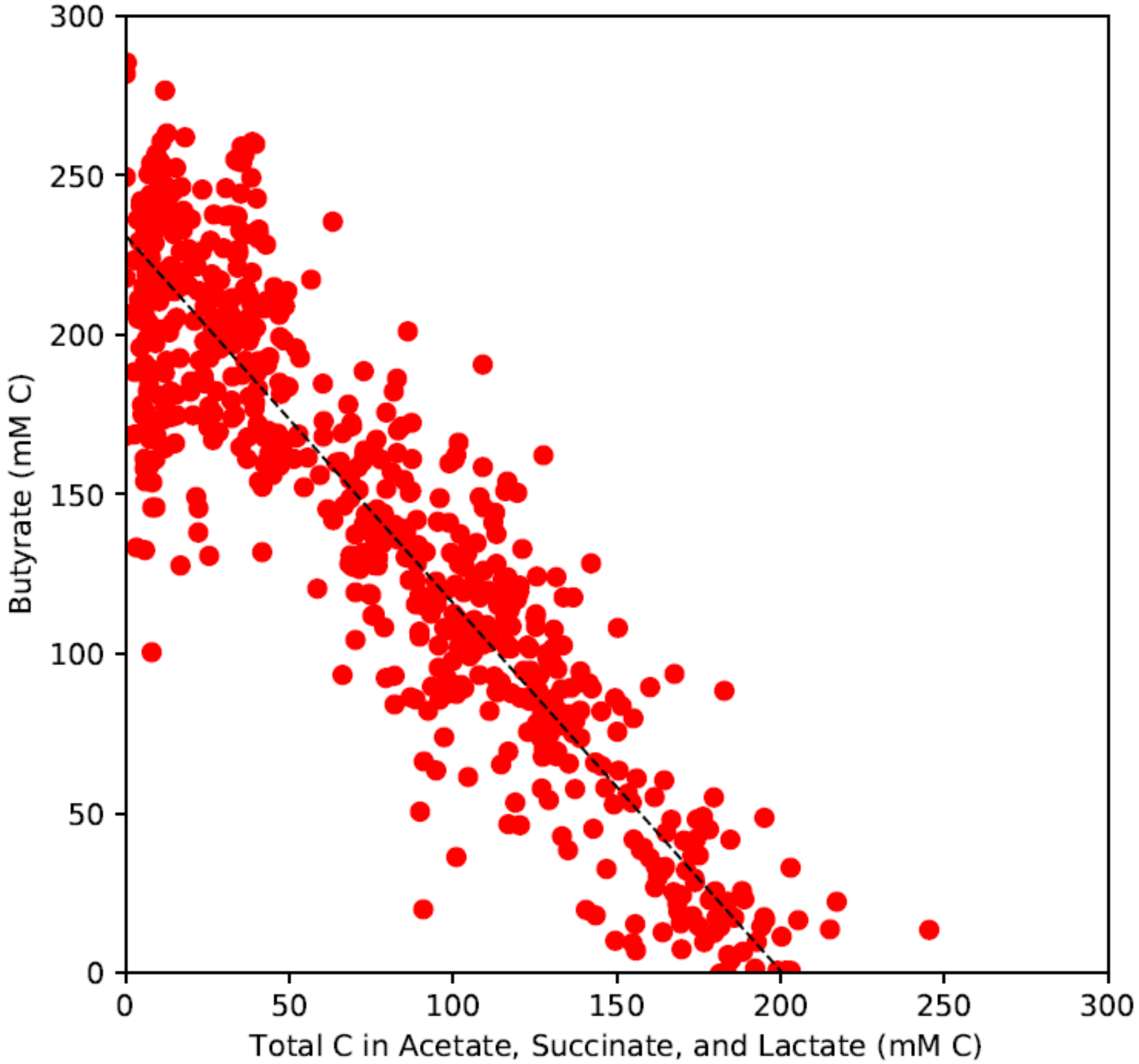


**Figure S4. Trade-off in organic acid production.** Scatter plot of total carbon in acetate, succinate, and lactate versus carbon in butyrate for communities with greater than 10 species. Each point indicates a biological replicate of a community. Dashed line indicates the linear regression ($y=-1.15x+231$, $r=-0.93$, $p=7*10^{-293}$).

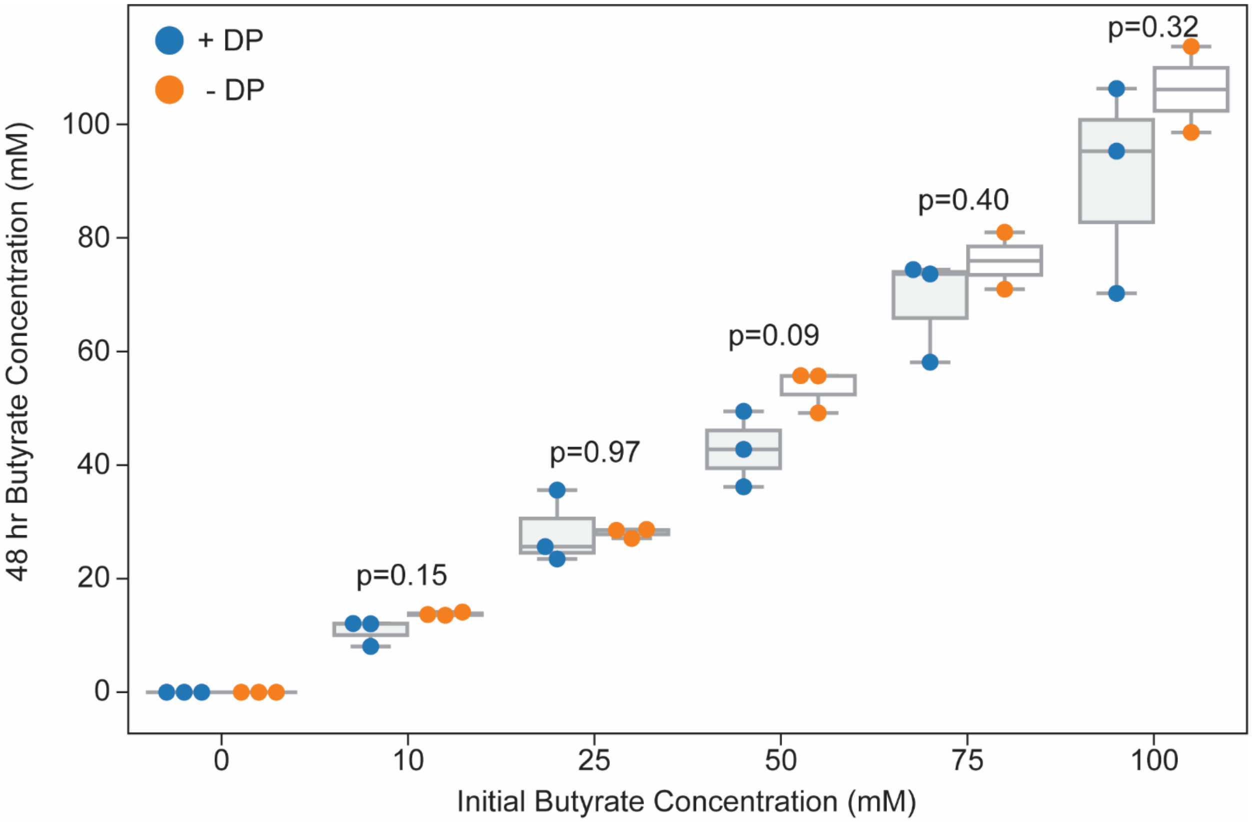


**Figure S5. Butyrate concentration in the presence and absence of DP.** Categorial scatter plot of butyrate concentrations in the presence of DP cultured for 48 hours in media supplemented with the indicated concentration of sodium butyrate (blue) or in a media control incubated for the same period of time (orange). The p-values were computed using a student's t-test (unequal variance).

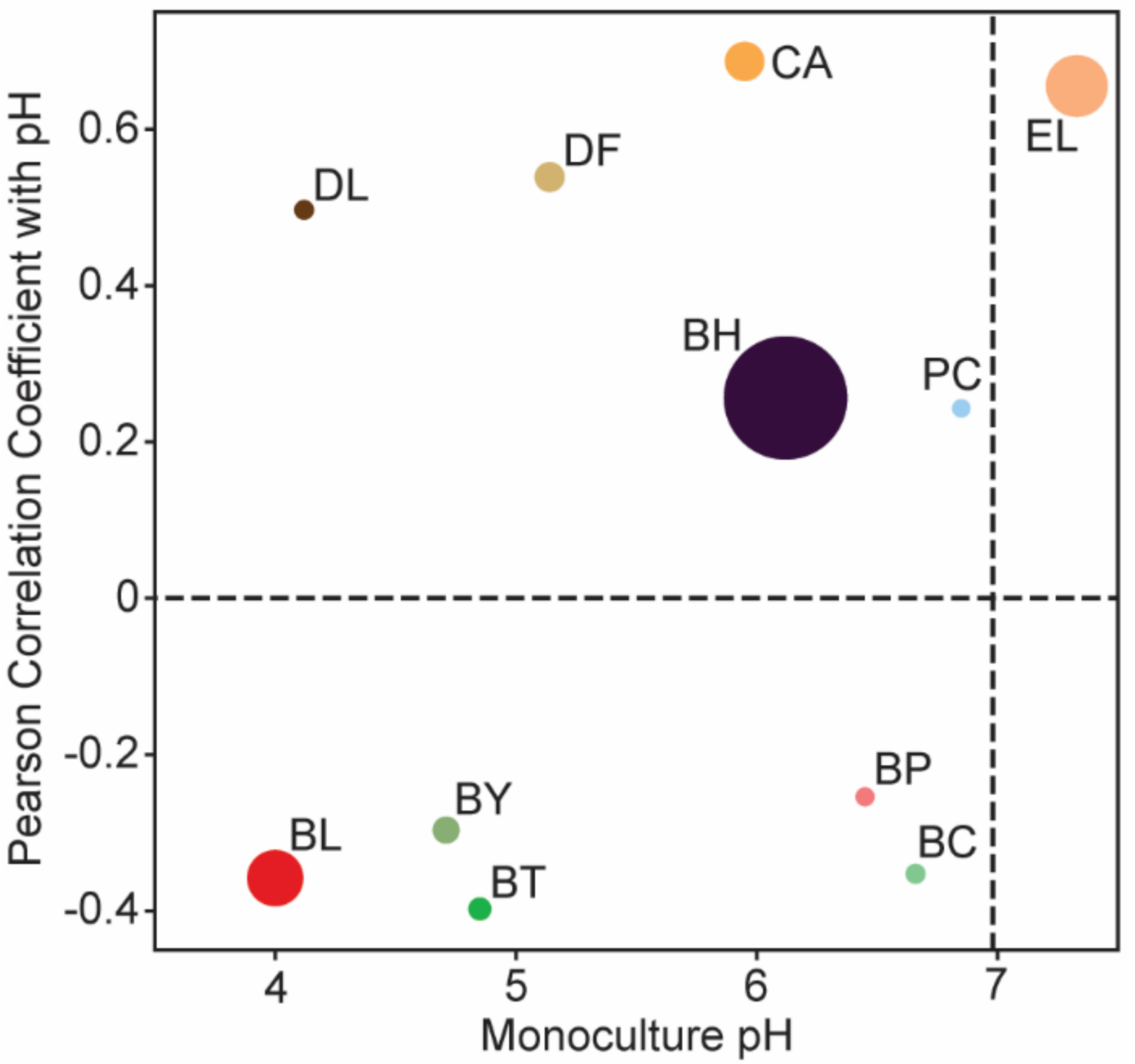


**Figure S6. Relationship between environmental pH of monocultures and correlation between species abundance and environmental pH.** Scatter plot of environmental pH of monospecies cultures at 48 hours versus the Pearson correlation coefficient of absolute species abundance with environmental pH at 48 hr in communities containing greater than 10 species. The size of the data points corresponds to the magnitude of the slope of the linear regression. Only those species with a statistically significant linear correlation ($p<0.05$) are shown. Vertical dashed line indicates the initial media pH.

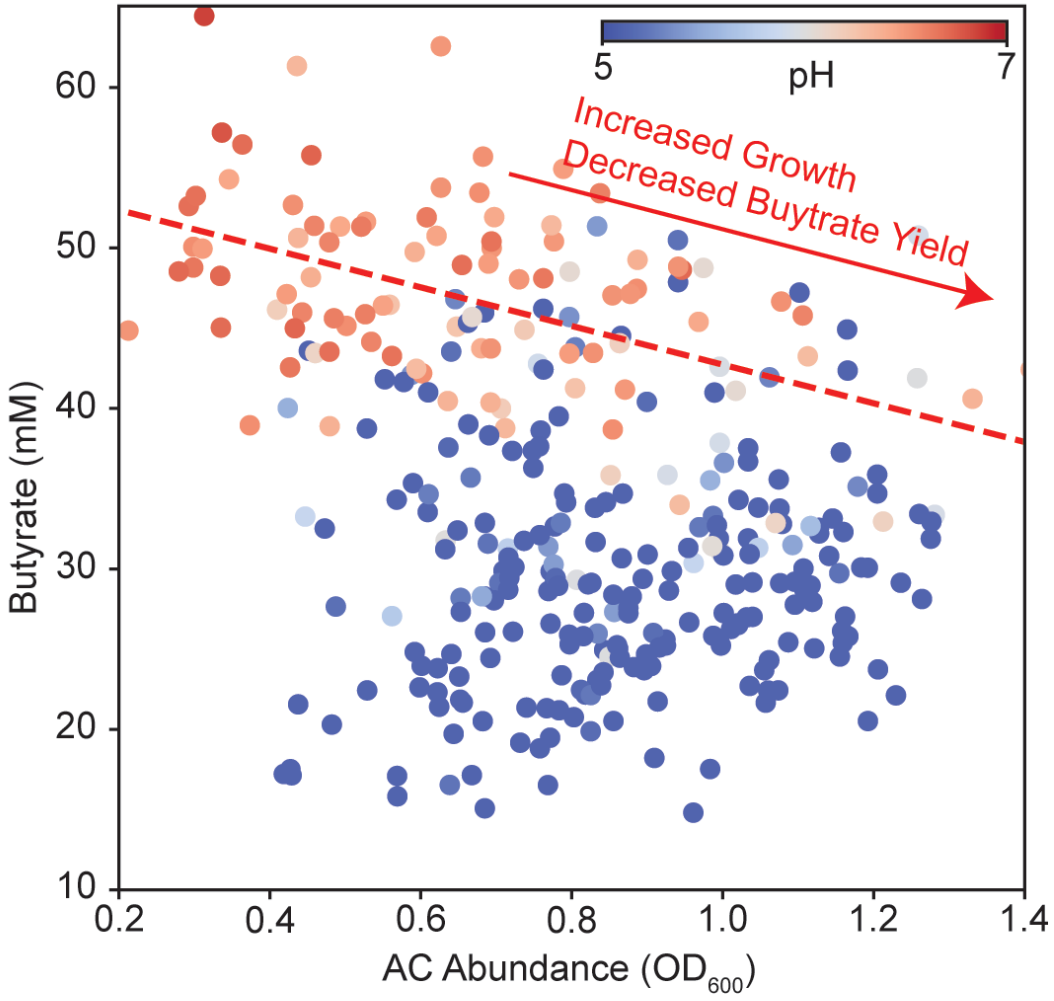


**Figure S7. Relationship between the abundance of AC and butyrate concentration.** Scatter plot of butyrate concentration versus the abundance of AC in communities from **a** containing AC (leave-one-out communities excluded). Each data point denotes a biological replicate of a community. Dashed line indicates the linear regression for data points with pH>6 (y=-12x+55, r=-0.45, p=3*$10^{-6}$). Linear regression for data points with pH<6 was not statistically significant (p=0.13).
